# Unique Proteomes Implicate Functional Specialization across Heterocysts, Akinetes, and Vegetative Cells in *Anabaena cylindrica*

**DOI:** 10.1101/2020.06.29.176149

**Authors:** Yeyan Qiu, Liping Gu, Volker Brözel, Douglas Whitten, Michael Hildreth, Ruanbao Zhou

## Abstract

In response to environmental changes, vegetative cells of *Anabaena cylindrica* can differentiate into two other cell types: a heterocyst for oxic N_2_-fixation, and an enlarged spore called akinete for stress survival. Akinetes normally differentiate from vegetative cells adjacent to heterocysts. Heterocysts inhibit nearby cells from differentiating into heterocysts but can induce adjacent cells to become akinetes, a rare embryogenetic induction in prokaryotes. The mechanism for a patterned differentiation in *A. cylindrica* has been little studied. Here, we isolated three types of cells from *A. cylindrica* to identify their proteomes using LC-MS/MS.

A total of 1395 proteins were identified, including 664 proteins from akinetes, 751 proteins from heterocysts, and 1236 proteins from vegetative cells. There were 45 proteins (33 novel proteins) found exclusive to akinetes, 57 heterocyst-specific proteins (33 novel proteins), including *nif* gene products, and 485 proteins exclusively in vegetative cells. Our proteomic data suggest that akinetes, unlike the typical spores of bacteria, perform unique biochemical functions that collaborate with both heterocysts and vegetative cells. A HAVe model for collaboration among heterocysts, akinetes and vegetative cells is proposed to illustrate the metabolic network of cyanophycin and carbohydrates based on the distribution of their biosynthesis related proteins in three types of cells. Interestingly, cell division proteins, DNA replication proteins, some carboxysomal proteins including RuBisCO and proteins in photosystems I, II were found abundant in heterocysts, the non-dividing cells dedicated exclusively to oxic N_2_-fixation. The identification of the akinete and heterocyst proteomes enables the pursuit of genetic studies into the patterned differentiation of akinetes and heterocysts.

## Introduction

Cyanobacteria are the only prokaryotes capable of oxygenic photosynthesis (Gantt, 2011), and are widely believed to be the ancestors of chloroplasts (Martin et al., 2002). Many cyanobacteria are also capable of photosynthetic fixing atmospheric dinitrogen (N2) (Kumar et al., 2010a). While some cyanobacteria follow a single-cell lifestyle, multicellularity in this group first evolved 2.5 billion years ago (Schirrmeister et al., 2011). Many cyanobacteria are capable of complex biochemical transformations in response to different physicochemical environments. Photosynthesis occurs in light and yields oxygen while N_2_ fixation requires a highly reduced environment (Kumar et al., 2010b). Unicellular cyanobacteria such as *Cyanothece sp*. ATCC 51142 solve this through a circadian clock to separate photosynthesis and N_2_ fixation temporarily into light and dark periods (Cerveny et al., 2013). Spatial division of labor in multicellular cyanobacteria appears more efficient at energy capture than temporal separation as occurs in unicellular cyanobacteria (Rossetti et al., 2010). Some filamentous cyanobacteria can differentiate to form four cell types: photosynthetic vegetative cells, N_2_-fixing heterocysts, akinetes, and small motile filaments called hormogonia (Rippka and Herdman, 1985; Flores and Herrero, 2010b). Akinetes developed from vegetative cells but are capable of germinating to produce young vegetative cells. Heterocysts develop from vegetative cells to form terminally differentiated, non-dividing cells functionally specialized for oxic N_2_-fixation. Heterocysts are formed in filamentous cyanobacteria in response to the depletion of fixed nitrogen (Mitschke et al., 2011). They develop every 10 to 20 cells along the filament (Kumar et al., 2010b), and are larger and more round than vegetative cells. The cell envelope of heterocysts is thicker, and with two additional envelope layers: heterocyst-specific glycolipids (HGL) and an outer polysaccharide layer (HEP). These extra two envelope layers impede the entry of oxygen to protect nitrogenase in the heterocysts (Flores and Herrero, 2010a). Heterocysts have diminished levels of pigments, and photosystem II is degraded to shut down O_2_-producing reactions (Thomas, 1970; Donze et al., 1972). Thus, the heterocyst creates a micro-oxic environment to house the oxygen-sensitive nitrogenase. However, photosystem I (PS I) is kept intact to generate ATP using light energy for N_2_-fixation through cyclic photophosphorylation (Wolk and Simon, 1969; Tel-Or and Stewart, 1976). Theretofore, nitrogen fixation in heterocysts is a uniquely solar-powered process, which is distinct from N_2_-fixation by any other N_2_-fixing bacteria. The wall between vegetative cells and heterocysts contains intercellular channels called septosomes, which allow for exchange of metabolites. Reductants such as sucrose and fixed carbon are obtained from vegetative cells, while heterocysts fix N_2_ and provide amino acids to the vegetative cells in a filament (Thomas et al., 1977; Muro-Pastor and Hess, 2012).

Some cyanobacteria can form akinetes, spore-like cells resistant to desiccation and freezing temperatures, that are able to germinate into new vegetative cells under favorable conditions (Perez et al., 2015). Unlike endospores of *Bacillus*, akinetes are susceptible to heat and long-term exposure to vacuum (Olsson-Francis et al., 2009). Akinetes are larger than vegetative cells (Singh and Montgomery, 2011) and contain large quantities of reserve products, mainly glycogen (Sarma et al., 2004) and the nitrogen storage polypeptide polymer cyanophycin (Sukenik et al., 2015). Akinetes are enveloped in a thick protective coat (Meeks et al., 2002). They begin to differentiate from vegetative cells during the late exponential phase of growth. Increasing culture density and decreasing light penetration accelerate the formation of akinetes. Intriguingly akinetes normally form adjacent to heterocysts in *Anabaena cylindrica* (Figure 1A), implying that these akinetes may play a role in transportation of N and C between vegetative cells and heterocysts besides their survival role in stress conditions. The significant morphological and metabolic changes observed in heterocysts and akinetes suggest unique phenotypes underpinned by complex regulatory pathways.

**FIGURE 1.**
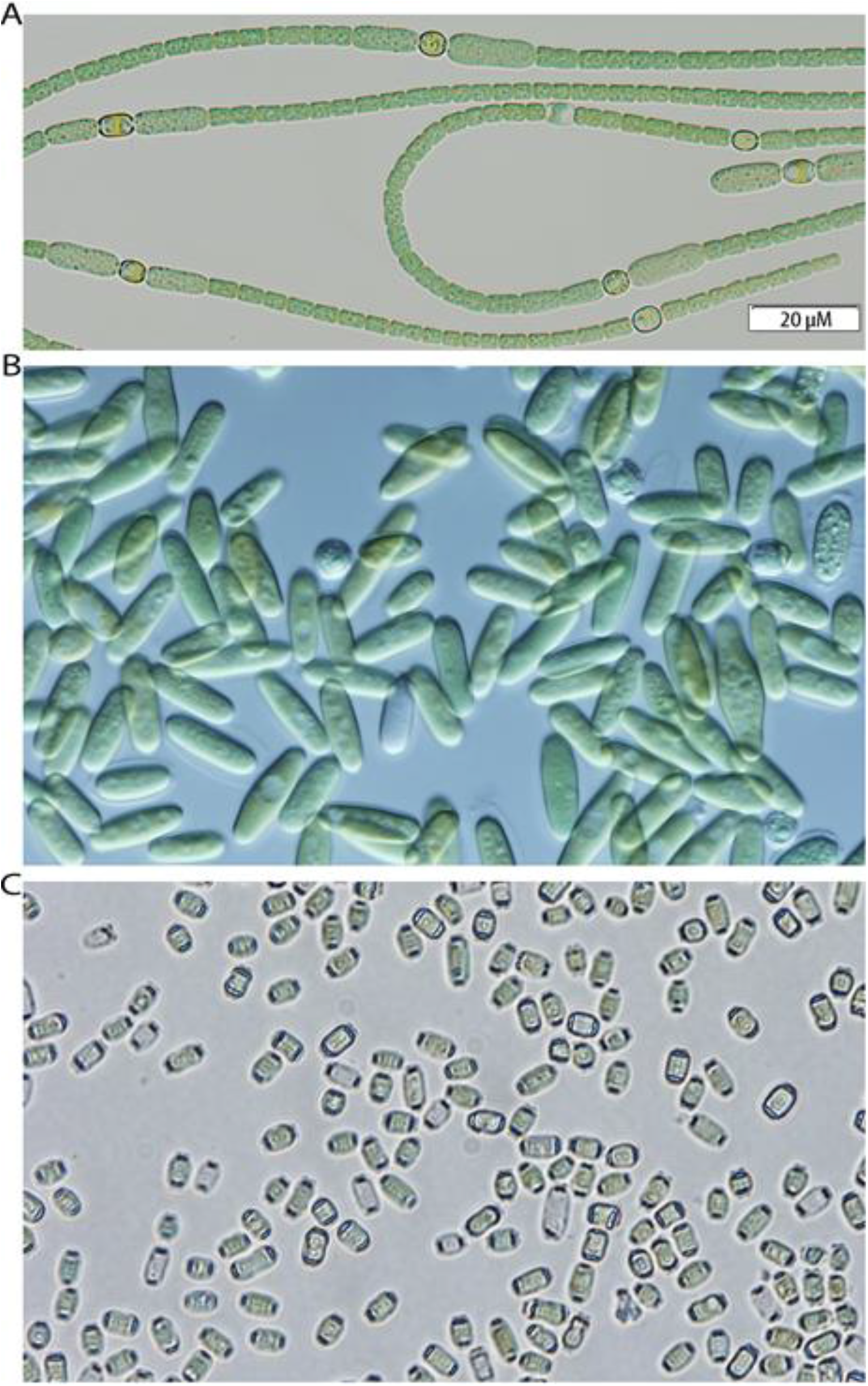
*Anabaena cylindrica* ATCC 29414 has three types of cells in filaments (A). The purity of the isolated akinetes (B) and heterocysts (C) was analyzed by differential interference contrast microscopy. **A:** Akinetes or developing akinetes, **H:** Heterocyst, **V:** Vegetative cell. Scale bar for panel A-C is 20 μm. The purity of the heterocysts and akinetes was 99.52 ± 0.48% and 96.17 ± 0.72%, respectively.

Many genes have been reported to be involved in regulating heterocyst differentiation. HetR is a master regulator specifically required for heterocyst differentiation (Buikema and Haselkorn, 1991; Zhou et al., 1998; Huang et al., 2004). Several regulatory genes *nrrA* (Ehira and Ohmori, 2011), *ccbP* (Hu et al., 2011), *hetN* (Higa et al., 2012), *hetF, patA* (Risser and Callahan, 2008), *patN* (Risser et al., 2012), *patU* (Meeks et al., 2002), *hetZ* (Zhang et al., 2007), *patS* (Yoon and Golden, 1998; Hu et al., 2015) and *hetP* (Videau et al., 2016) were also found to play very important roles during heterocyst differentiation and its pattern formation. The heterocyst-specific NsiR1 small RNA was recently discovered as an early marker in this process (Muro-Pastor, 2014). Although these genes are clearly involved in the regulation of heterocyst development, their biochemical functions remain to be determined. Unfortunately, the genetic regulation of akinete formation is completely unknown. So far, the only reported akinete-specific protein is AvaK from *Anabaena variabilis* (Zhou and Wolk, 2002). There has been no proteomic study for akinetes to date although a quantitative shotgun proteomics study of heterocysts was reported for *Anabaena sp*. PCC 7120 (Ow et al., 2008; Pandey et al., 2012; Agrawal et al., 2014; Panda et al., 2014) and *Nostoc punctiforme* (Liang et al., 2012; Sandh et al., 2014).

*A. cylindrica* can form N_2_-fixing heterocysts under both depleted and replete nitrate conditions (Meeks et al., 1983), which is different from other heterocyst forming cyanobacteria, such as *Anabaena* sp. strain PCC 7120 (Borthakur and Haselkorn, 1989), *Anabaena variabilis* (Thiel et al., 1995) and *Nostoc punctiforme* (Summers and Meeks, 1996), whose vegetative cells can differentiate into heterocysts only in response to deprivation of combined nitrogen. Moreover, vegetative cells of *A. cylindrica* can also differentiate into akinetes (arrowheads labeled A), spore-like cells for stress survival. Akinetes (15 ~20μm in length) are about 10 times larger than vegetative cells, and normally develop adjacent to heterocysts within the same filament (Figure 1A), providing a rare opportunity to elucidate what appears to be an embryogenetic induction in a prokaryote (Wolk, 1966). Unfortunately, the differentiation of akinetes, heterocysts as well as akinete juxtaposition to heterocysts have not heretofore been studied genetically due to the lack of a genetic transformation method for this organism.

We sought to characterize the phenotype of akinetes of *A. cylindrica* through proteomic analysis, contrasting it to the phenotype of heterocysts and vegetative cells. *A. cylindrica* ATCC 29414 was selected for this study because it differentiates readily into both heterocysts and akinetes in dilute Allen and Arnon medium (AA/8) without combined nitrogen (Hu et al., 1981). Its akinetes are large and readily separated from heterocysts and vegetative cells. Our proteomic data suggest that akinetes, unlike the typical spores of bacteria, perform unique biochemical functions that collaborate with both heterocysts and vegetative cells.

## Material and Methods

### Isolation of akinetes and heterocysts

Isolation of akinetes and heterocysts was based upon the CsCl density gradient centrifugation (Wolk, 1968) with the following modification. Briefly *A. cylindrica* ATCC 29414 was grown in nitrate free AA/8 medium under continuous light (60 μE/m^2^/s, 150 rpm, 30°C) for 30 days (OD_700_≈0.15) to allow heterocyst and akinete development. Cultures were harvested (6,400 × *g* 15 min, 4°C), resuspended in ddH_2_O, and the vegetative cells were disrupted by passing the suspension through a Nano DeBEE-30 high pressure homogenizer (BEE International) at 4,500 psi and then at 5,000 psi. Akinetes and heterocysts were sedimented (4,000 × *g* 10 min, and 4°C) and washed four times with ddH_2_O to remove the vegetative cell debris. There were two distinct layers formed in the last wash pellet. The upper layer was suspended in 1.55 g/mL CsCl density solution, and transferred into in an ultracentrifugation tube. The bottom layer was suspended in 1.45 g/mL CsCl and carefully transferred on-top of the upper layer suspension in the same ultracentrifugation tube. Two distinct fractions were collected from the first CsCl density gradient centrifugation (17,000 × *g*, 60 min, 4°C in a fixed angle MLA-55 rotor, Beckman Coulter), each were sedimented (4000 × *g*, 30 min), and washed with 3x ddH_2_O. The heavy fraction was suspended in 1.45 g/ml CsCl solution, and re-centrifuged as before. The light fractions from the first and second centrifugations were pooled, suspended in 1.45 g/ml CsCl solution, and re-centrifuged. The supernatant fraction from this third centrifugation was suspended in 1.3 g/ml CsCl solution and re-centrifuged. The resultant pellets from the second and third centrifugations (containing highly purified akinetes) and pellet from the 4^th^ centrifugation (containing highly purified heterocysts) were washed with ddH_2_O as above. The purity of the heterocysts (99.52 ± 0.48%) and akinetes (96.17 ± 0.72%) was examined by differential interference contrast microscopy (AX70 upright, Olympus).

### Total protein extraction and SDS-PAGE purification

The purified heterocysts or akinetes were suspended in Phosphate Buffered Saline (PBS) containing 1% N-lauroyl sarcosine and protease Inhibitors [Complete, Mini Protease Inhibitor Cocktail Tablets (Roche)] at 1 tablet per 10 mL]. Cells were disrupted on ice by ultra-sonication (Branson digital sonifier 450) for 12 x 5 s bursts with 15 s interval at an amplitude of 60%. Cell lysates were collected (13,000 × *g* 20 min) respectively. To extract total proteins from the vegetative cells, *A. cylindrica* ATCC 29414 cells grown in AA/8N (Hu et al., 1981) for 8 days (OD_700_≈0.042) were harvested (6,400 × *g*, 15 min, 4°C) and resuspended in PBS buffer containing 1% N-lauroyl sarcosine and protease inhibitors. Vegetative-cell lysate was obtained by passing the cell suspension through a Nano DeBEE-30 High Pressure Homogenizer (BEE International) once at 1,000 psi (only the vegetative cells were disrupted at this pressure), and removing unbroken cells via centrifugation (4,000 × *g*, 10 min, 4°C).

Total proteins from each type of cell lysate were precipitated with 10% trichloroacetic acid (TCA) overnight at 4°C, sedimented (16,000 × *g*, 30 min), and washed three times with 80% methanol and three times with 80% acetone. The pellets were resuspended in sodium dodecyl sulfate-polyacrylamide gel electrophoresis (SDS-PAGE) loading buffer containing 1% N-lauroyl sarcosine, boiled for 3 min, clarified by centrifugation at 16,000 × *g* for 20 min at 25°C, and subjected to a 12% SDS-PAGE (Bio-Rad Mini-PROTEAN^®^ Comb, 5-well, 1.0 mm) at 200 mV for approximately 15 min, until all the proteins just entered the resolving gel. The gel was stained with Coomassie Brilliant Blue R-250 for band excision and analysis (Supplementary Figure S1).

### In-gel tryptic digestion and protein identification by LC-MS/MS

The protein gel bands (Supplementary Figure S1) were excised and in-gel tryptic digestion was performed according to Shevchenko (Shevchenko et al., 1996) with the following modifications. Briefly, gel slices were dehydrated with acetonitrile (ACN) for approximately 5 min incubation and repeated this process until they appear to shrink in size and show a chalk white color. The time required and number of washes vary with gel size and composition. The chalk white color gel was then incubated with 100 mM ammonium bicarbonate (NH_4_HCO_3_) containing 10 mM dithiothreitol (DTT, pH ≈ 8.0) for 45 min at 56°C, dehydrated again and incubated with 100 mM NH_4_HCO_3_ containing 50 mM iodoacetamide for 20 min in the dark, and then washed with 100 mM NH_4_HCO_3_ and dehydrated again. Approximately 50 μL trypsin solution (0.01 μg/μL sequencing grade modified trypsin (Promega, #V5111) in 50 mM NH_4_HCO_3_) was added to each gel slice so that the gel was completely submerged, and then incubated at 37°C for overnight. The tryptic peptides were extracted with 60% ACN/1% TCA from the gel by water bath sonication (Aquasonic 150T sonicating water bath which puts out 135W. Sonication is done 2 x 20s) and concentrated in a SpeedVac to 2 μL.

For heterocyst and akinete samples, the extracted peptides were re-suspended in 20 μL 2% ACN/0.1% trifluoroacetic acid (TFA), 10 μL were injected by a nanoAcquity Sample Manager and loaded for 5 min onto a Symmetry C18 peptide trap (5 μm, 180 μm x 20 mm) (Waters) at 4 μL/min in 2% ACN/0.1% Formic Acid. The bound peptides were eluted onto a BH130 C18 column (1.7 μm, 150 μm x 100 mm, Waters) using a nanoAcquity UPLC (Waters) (Buffer A = 99.9% Water/0.1% Formic Acid, Buffer B = 99.9% Acetonitrile/0.1% Formic Acid) with a gradient of 5% B to 30% B over 228 min, ramping to 90% B at 229 min and holding for 1 min, and then ramping back to 5% B at 231 min, and holding for equilibration prior to the next injection for a total run time of 240 min. The eluted peptides were sprayed into a LTQ-FT-ICR Ultra hybrid mass Spectrometer (Thermo Scientific) using an ADVANCE nanospray source (Bruker-Michrom). Survey scans were taken in the FT (25,000 resolution determined at *m/z* 400) and the top five ions in each survey scan were then subjected to automatic low energy collision induced dissociation (CID) in the LTQ.

For the vegetative-cell sample, 5 μL of the extracted peptide suspension was injected (to the sample loop which is then backflushed using solvent A directly to the column) by EASYnLC and the peptides separated through an Acclaim PepMap RSLC column (0.075 mm x 150 mm C18, Thermo Scientific) with the same gradient as above. The eluted peptides were sprayed into a Q Exactive hybrid quadrupole-Orbitrap mass spectrometer using a Nanospray Flex™ Ion Sources (Thermo Scientific). Survey scans were taken in the Orbi trap (35,000 resolution determined at *m/z* 200) and the top ten ions in each survey scan were then subjected to automatic higher energy collision induced dissociation (HCD) with fragment spectra acquired at 17,500 resolution (by convention this is a dimensionless measurement).

For protein identification, the resulting MS/MS spectra were converted to peak lists using Mascot Distiller, v2.5.1.0 (www.matrixscience.com) and searched against a protein sequence database containing *A. cylindrica* ATCC 29414 (http://scorpius.ucdavis.edu/gmod/cgi-bin/site/anabaena02?page=gblast), *A. cylindrica* PCC 7122 entries (http://www.ncbi.nlm.nih.gov/genome/?term=anabaena+cylindrica+7122+genome) and common laboratory contaminants downloaded from www.thegpm.org. All searches were performed using the Mascot searching algorithm, v 2.4. The Mascot output was then analyzed using Scaffold, v4.3.4 (www.proteomesoftware.com) to probabilistically validate protein identifications at 1% FDR. The quantification value was calculated using Normalized Total Spectra (For details, see Supplementary Materials). The mass spectrometry proteomics data have been deposited to the ProteomeXchange 213 Consortium via the PRIDE (Vizcaino et al., 2016) partner repository with the dataset identifier PXD006041.

## Results

### Proteomic analysis of heterocysts, akinetes, and vegetative cells

To unlock the cellular function of akinetes and the protein network among akinetes, heterocysts, and vegetative cells in *A. cylindrica* ATCC 29414, we performed proteomics through LC-MS/MS. A total of 12616 tryptic peptides were collected and 1426 proteins were identified, including 1395 ORF proteins from *A. cylindrica* ATCC 29414, 14 proteins from common laboratory contaminants, 14 decoy proteins for determination of the false discovery rate, and 3 ORFs (Anacy_0074, Anacy_3940 and Anacy_5216) in *A. cylindrica* PCC 7122 matched to intergenic regions of *A. cylindrica* ATCC 29414 genome (Supplementary Table S2).

Our LC-MS/MS proteomics analysis identified 664 proteins from akinetes, 751 from heterocysts, and 1236 from vegetative cells, with 448 proteins common to all three cell types. There were 45 akinete-specific (Supplementary Table S1), 57 heterocyst-specific (Supplementary Table S1), and 485 vegetative cell-specific proteins (Figure 2, Supplementary Table S2). Interestingly, phycocyanin alpha (ORF: 3613) and beta (ORF: 3614) subunits, allophycocyanin beta subunit (ORF: 1908), phycobilisome protein (ORF: 1909), beta subunit of mitochondrial ATP synthase (ORF: 3788), translation elongation factor 1A (EF-1A/EF-Tu, ORF: 5853) and ribulose 1,5-bisphosphate carboxylase (RuBisCO) large (ORF: 6007) and small subunit (ORF: 6009) were among most abundant proteins in all three cell types.

**FIGURE 2.**
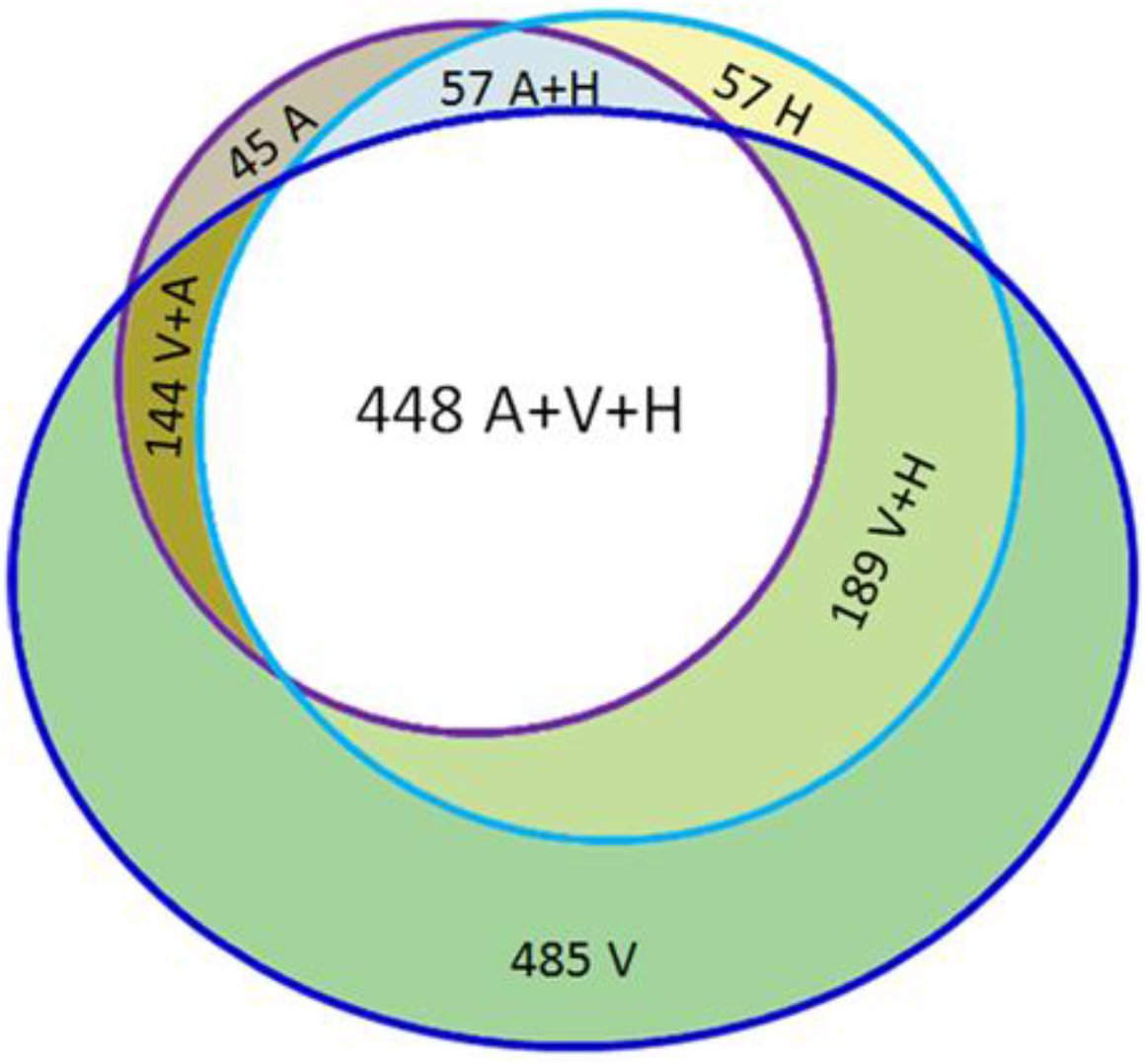
Venn analysis showing the proteomic profiles from akinetes (A), heterocysts (H), and vegetative cells (V). **45 A:** 45 proteins were detected exclusive to akinetes; **57 H:** 57 proteins were found to be heterocyst-specific; **485 V:** 485 proteins were found exclusively in vegetative cells; **57 A+H:** 57 proteins were detected in both akinetes and heterocysts; **144 V+A:** 144 proteins were detected in both vegetative cells and akinetes; **189 V+H:** 189 proteins were detected in both vegetative cells and heterocysts; and **448 A+V+H:** 448 proteins were found to be common to all three cell types.

### Nitrogen fixation in heterocysts

The *nif* genes encode subunits of nitrogenase for reducing atmospheric N_2_ to ammonia. Other heterocyst-specific genes encode proteins involved in regulating heterocyst development and N_2_ fixation, and inactivation of these genes showed diminished or ceased diazotrophic growth in the presence of oxygen due to impaired nitrogenase activity, or forming no or dysfunctional heterocysts (Lechno-Yossef et al., 2011). The above genes are collectively called ‘FOX’ genes (incapable of N_2_-fixation in the presence of oxygen) (Lechno-Yossef et al., 2011). LC-MS/MS identified 27 FOX proteins (Table 1) and 57 heterocyst-specific proteins (Supplementary Table S1). Heterocysts had 19 Fox proteins but eleven were also found in akinetes (Table 1).

**TABLE 1.** Distribution of FOX proteins, photosystem I and II proteins, and akinete marker protein in akinetes, heterocysts, and vegetative cells.

| ORF# | Annotation | Akinete | Heterocyst | Veget. Cell |
| --- | --- | --- | --- | --- |
|  |  | Normalized quantitative value |  |  |
| FOX genes |  |  |  |  |
| 2715 | Histidine kinase HepN | 0 | 0 | 0.45 |
| 4573 | Neutral invertase InvB | 0 | 0 | 0.45 |
| 4916 | Hypothetical protein Npun_R1723, FraG/SepJ | 0 | 0 | 0.45 |
| 6434 | Protein serine/threonine phosphatase PrpJ1 | 0 | 0 | 0.45 |
| 222 | Nitrogen-fixing NifU-like protein | 0 | 0 | 0.9 |
| 1433 | Response regulator receiver protein DevR | 0 | 0 | 0.9 |
| 2881 | ParB-like partition protein, HGL region containing | 0 | 0 | 1.8 |
| 3957 | Processing proteinase Abp2 | 0 | 0 | 2.71 |
| 3514 | Polyketide-type polyunsaturated fatty acid synthase PfaA/HglF | 0 | 2.45 | 0 |
| 5142 | FHA domain containing protein FraH | 0 | 2.45 | 0.45 |
| 4208 | FHA modulated glycosyl transferase/transpeptidase PBP3 | 2.66 | 2.45 | 0.45 |
| 3826 | Glucose-6-phosphate 1-dehydrogenase Zwl | 7.99 | 2.45 | 1.35 |
| 5437 | Nitrogenase molybdenum-iron protein alpha chain NifD | 10.66 | 4.9 | 0 |
| 907 | Nitrogenase molybdenum-iron cofactor biosynthesis protein NifN | 0 | 7.36 | 0 |
| 3521 | Glycosyl transferase (group 1) HglT | 2.66 | 7.36 | 0 |
| 1865 | Hypothetical protein Npun_F0815, Asp-, glu- rich product | 0 | 7.36 | 5.86 |
| 4777 | cytochrome c oxidase subunit II coxB2 | 0 | 7.36 | 0 |
| 1565 | Histidinol dehydrogenase HisD | 5.33 | 9.81 | 9.02 |
| 6242 | ABC transporter related DevA | 0 | 12.26 | 0 |
| 6485 | Histone-like DNA-binding protein HanA | 2.66 | 12.26 | 19.4 |
| 715 | Polyketide synthase thioester reductase subunit HglB | 0 | 14.71 | 0 |
| 6701 | Fe-S cluster assembly protein NifU | 0 | 22.07 | 0 |
| 3078 | cytochrome c oxidase subunit II coxB3 | 10.66 | 22.07 | 0 |
| 911 | Nitrogenase FeMo beta subunit protein NifK | 10.66 | 22.07 | 0.45 |
| 153 | Outer membrane efflux protein HgdD | 23.98 | 22.07 | 4.06 |
| 3764 | Hypothetical protein Npun_R5769 Abp3 | 37.3 | 34.33 | 11.73 |
| 3480 | Putative transcriptional regulator (Crp/Fnr family) DevH | 37.3 | 44.14 | 12.18 |
| 1651 | Polysaccharide export protein | 39.96 | 49.04 | 2.26 |
| 3002 | Nitrogenase iron protein NifH | 0 | 61.3 | 0.45 |
| Photosystems |  |  |  |  |
| 3318 | Photosystem I assembly BtpA | 0 | 2.45 | 0.45 |
| 5296 | Photosystem I assembly protein ycf3 | 0 | 0 | 1.35 |
| 1589 | Photosystem I assembly Ycf4 | 2.66 | 4.9 | 4.06 |
| 4482 | Photosystem I P700 apoprotein A2 | 50.62 | 17.17 | 12.18 |
| 5755 | Photosystem I iron-sulfur center | 18.65 | 26.97 | 18.04 |
| 2498 | Photosystem I reaction center protein PsaF, subunit III | 13.32 | 12.26 | 44.66 |
| 2501 | Photosystem I reaction center subunit XI | 45.29 | 122.6 | 59.09 |
| 1809 | Photosystem I reaction center subunit IV | 5.33 | 29.43 | 74.88 |
| 833 | Photosystem I reaction center subunit II | 39.96 | 100.54 | 163.3 |
| 1817 | Photosystem II reaction center Psb28 protein | 0 | 0 | 1.35 |
| 6255 | Photosystem II oxygen evolving complex protein PsbP | 5.33 | 4.9 | 4.96 |
| 3007 | Photosystem II protein D2 | 13.32 | 9.81 | 4.96 |
| 83 | Photosystem II reaction center protein H | 0 | 2.45 | 4.96 |
| 2046 | Photosystem q(b) protein | 5.33 | 9.81 | 7.22 |
| 1649 | Photosystem II 44 kDa subunit reaction center protein | 47.95 | 22.07 | 10.83 |
| 5644 | Photosystem II chlorophyll-binding protein CP47 | 47.95 | 29.43 | 14.89 |
| 3469 | Photosystem II 11 kDa protein | 21.31 | 22.07 | 17.59 |
| 3312 | Photosystem II oxygen evolving complex protein PsbU | 5.33 | 12.26 | 31.58 |
| 4725 | Photosystem II manganese-stabilizing protein PsbO | 42.62 | 71.11 | 104.66 |
| 755 | Photochlorophyllide reductase subunit N | 0 | 0 | 0.45 |
| **Akinete marker protein (AcaK43)** |  |  |  |  |
| 1647 | PRC-barrel domain-containing protein AvaK | 85.25 | 90.73 | 1.35 |

### Distinct distribution of Photosystem I and II proteins

PS I and PS II are well-known hallmarks primarily associated with the photosynthetic characteristics of vegetative cells. Of 20 photosystem proteins identified, all were present in vegetative cells, consistent with vegetative cells bearing both PS I and PS II proteins for fully functional photosynthesis and electron transfer (Table 1). The protochlorophyllide reductase subunit N catalyzing the penultimate step of chlorophyll biosynthesis (Yamazaki et al., 2006), PS I assembly protein Ycf3 (Wilde et al., 2001), and PS II reaction center Psb28 protein (Dobakova et al., 2009) were unique to vegetative cells of *A. cylindrica*. Notably, there were several PS I and PS II proteins in high abundance in both akinetes and heterocysts respectively (Table 1), e.g., PS I P700 apoprotein A2, PS II 44 kDa subunit reaction center protein, and PS II chlorophyll-binding protein CP47, suggesting that PS I and PS II may function partially in both heterocysts and akinetes.

### Akinete-specific protein AcaK43 (ORF: 1647) in *A. cylindrica*

The first reported akinete-specific protein was AvaK from *Anabaena variabilis* (Zhou and Wolk, 2002). AvaK homolog AcaK43 (ORF: 1647) was among the top 20 most abundant proteins in *A. cylindrica* akinetes (85.25 counts) and heterocysts (90.73 counts), while only trace amount was detected in vegetative cells, which is consistent with the previous report in *A. variabilis* that AvaK is an akinete marker protein (Zhou and Wolk, 2002). However, our proteomic data showed that AcaK43 was abundant in heterocysts of *A. cylindrica*. Fluorescence of GFP (green fluorescent protein) from *PacaK43-gfp* (promoter of *acaK43* fused to *gfp*) originates primarily in both akinetes and heterocysts of *A. cylindrica* (Zhou et al, unpublished observation). Furthermore, proteins homologous to AapN, Hap, Aet identified as akinete-specifically expressed genes using differential display at mRNA level in *Nostoc punctiforme* (Argueta et al., 2006) were below the limit of detection in our proteomic study. The presence of AcaK43 in both akinetes and heterocysts suggest that *A. cylindrica* is distinct from *A. variabilis* and *N. punctiforme* although they all are akinete/heterocyst-forming cyanobacteria.

### DNA/RNA/protein biosynthesis patterns among akinetes, heterocysts, and vegetative

LC-MS/MS identified a number of proteins related to nucleotide synthesis, DNA packing and repair, RNA and protein synthesis, and cell division within akinetes (Table 2). Nucleotide synthesis related protein (phosphoribosylformylglycinamidine synthase II) was found to be akinete-specific. Although DNA polymerases, single-strand binding protein (SSBP) and DEAD/DEAH box helicase domain-containing protein involved in DNA replication were not detected in akinetes, DNA gyrase subunit A and the DNA gyrase modulator Peptidase U62 were only found in akinetes. Akinetes have more chromosome copies per cell than in vegetative cells (Sukenik et al., 2012), so DNA gyrase might play an important role in DNA wrapping and packaging (Gore et al., 2006). Furthermore, DNA gyrase in akinetes may minimize the potential damage caused by light energy to these resting cells (Napoli et al., 2004). Moreover, a similar distribution pattern of DNA replication proteins was seen in heterocysts.

**TABLE 2.**
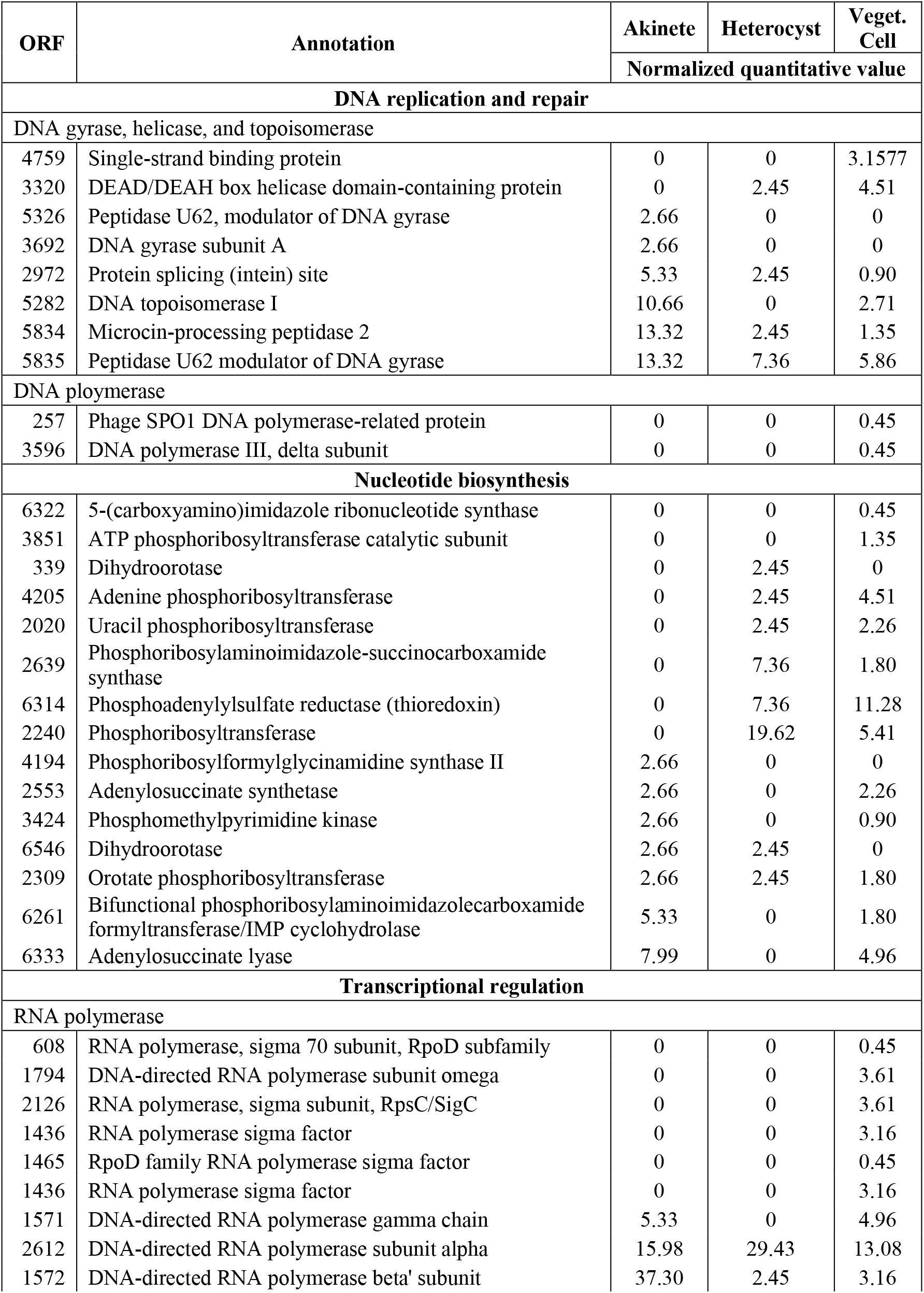
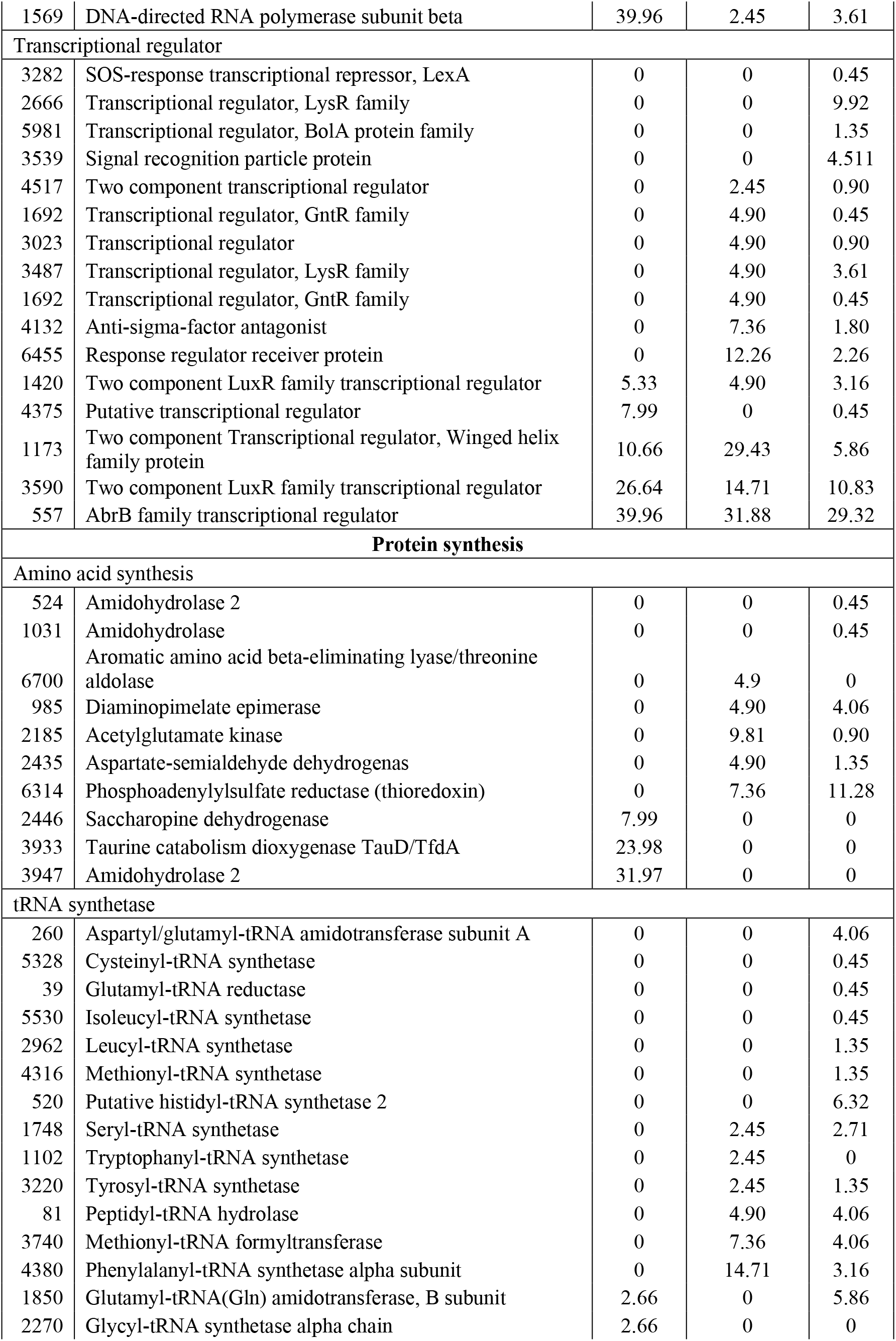
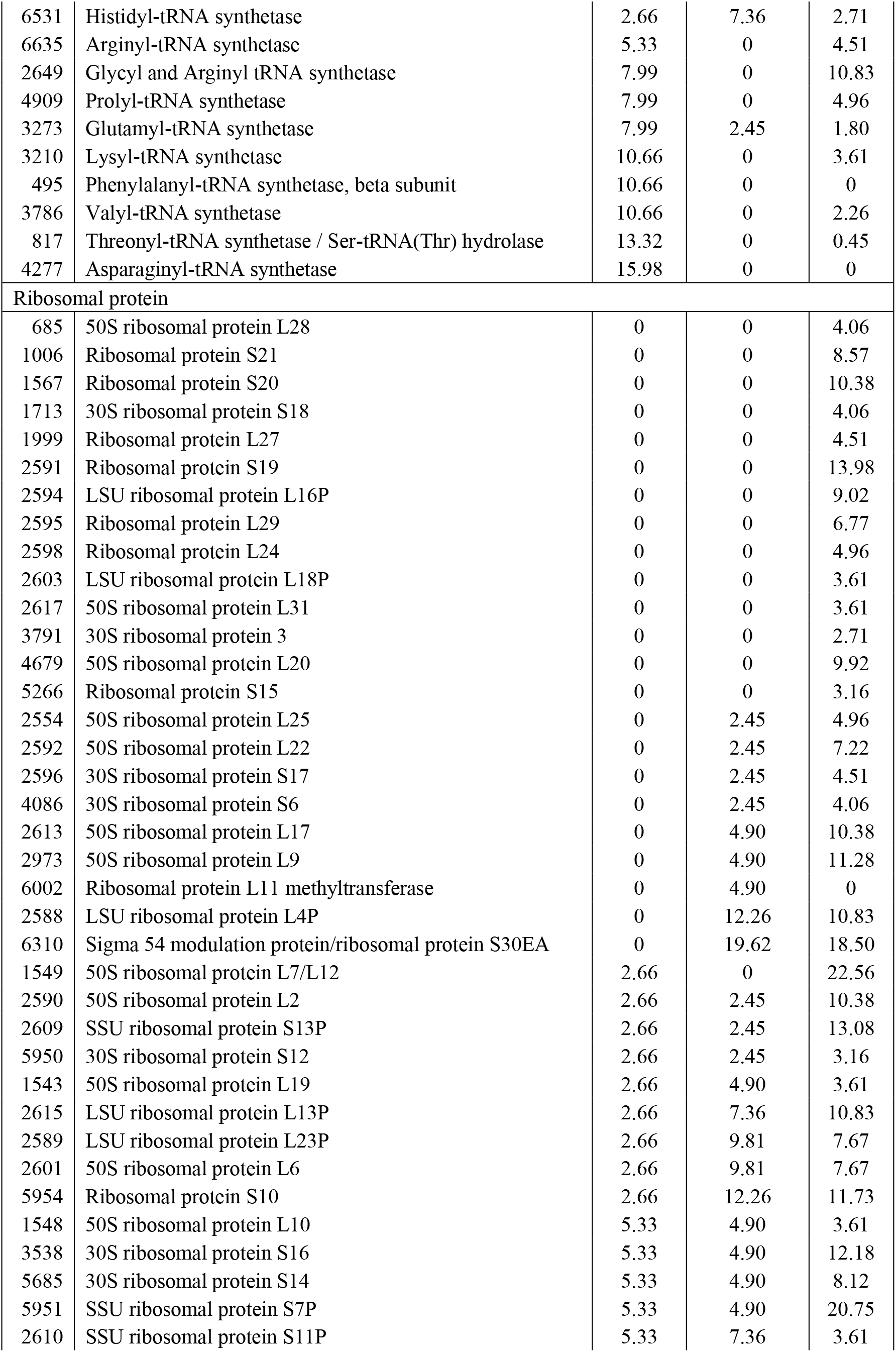
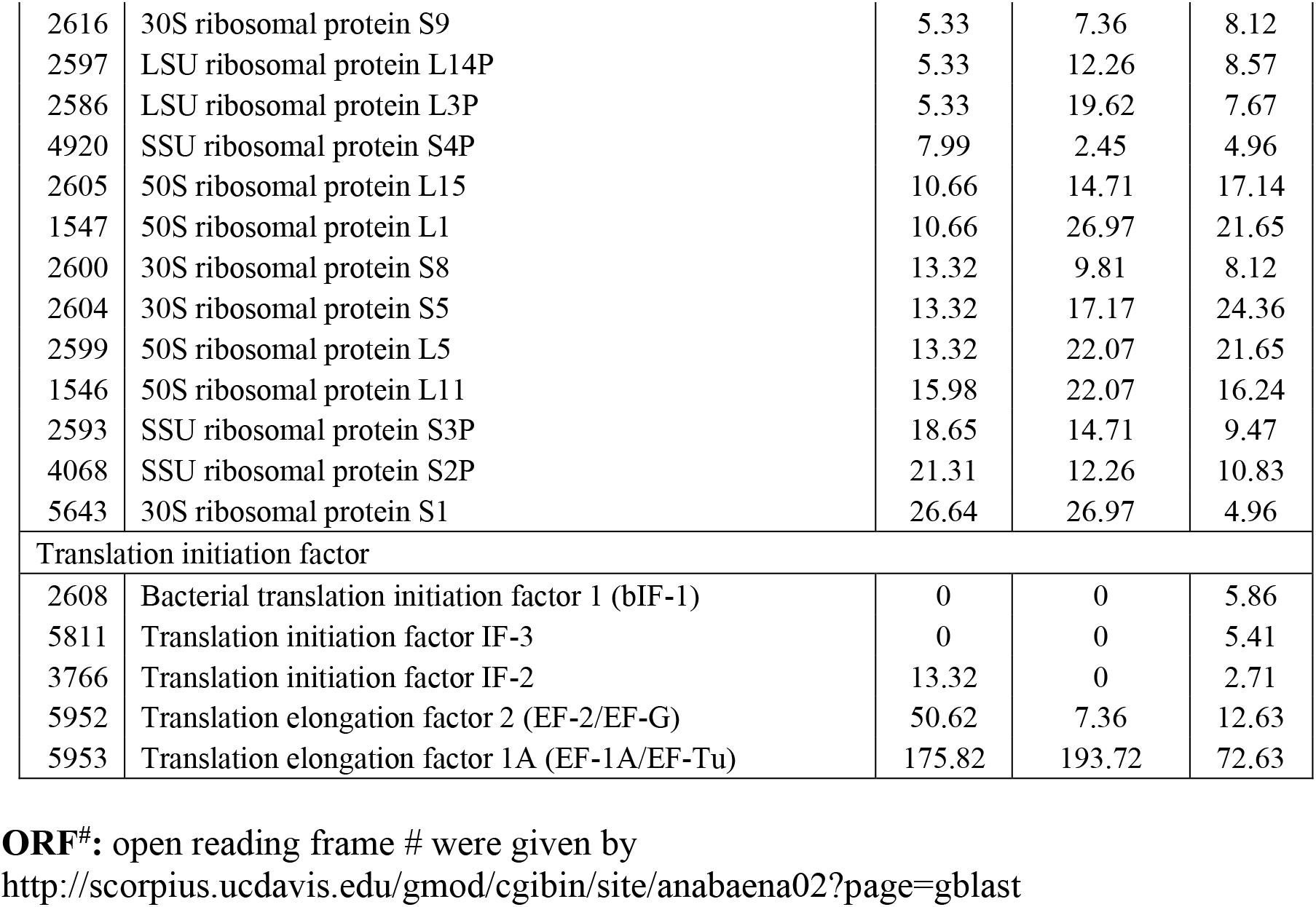
Proteins involved in biosynthesis of DNA, RNA, and protein, as well as cell division in akinetes, heterocysts, and vegetative cells respectively

| ORF | Annotation | Akinete | Heterocyst | Veget. Cell |
| --- | --- | --- | --- | --- |
|  |  | Normalized quantitative value |  |  |
| DNA replication and repair |  |  |  |  |
| DNA gyrase, helicase, and topoisomerase |  |  |  |  |
| 4759 | Single-strand binding protein | 0 | 0 | 3.1577 |
| 3320 | DEAD/DEAH box helicase domain-containing protein | 0 | 2.45 | 4.51 |
| 5326 | Peptidase U62, modulator of DNA gyrase | 2.66 | 0 | 0 |
| 3692 | DNA gyrase subunit A | 2.66 | 0 | 0 |
| 2972 | Protein splicing (intein) site | 5.33 | 2.45 | 0.90 |
| 5282 | DNA topoisomerase I | 10.66 | 0 | 2.71 |
| 5834 | Microcin-processing peptidase 2 | 13.32 | 2.45 | 1.35 |
| 5835 | Peptidase U62 modulator of DNA gyrase | 13.32 | 7.36 | 5.86 |
| DNA ploymerase |  |  |  |  |
| 257 | Phage SPO1 DNA polymerase-related protein | 0 | 0 | 0.45 |
| 3596 | DNA polymerase III, delta subunit | 0 | 0 | 0.45 |
| Nucleotide biosynthesis |  |  |  |  |
| 6322 | 5-(carboxyamino)imidazole ribonucleotide synthase | 0 | 0 | 0.45 |
| 3851 | ATP phosphoribosyltransferase catalytic subunit | 0 | 0 | 1.35 |
| 339 | Dihydroorotase | 0 | 2.45 | 0 |
| 4205 | Adenine phosphoribosyltransferase | 0 | 2.45 | 4.51 |
| 2020 | Uracil phosphoribosyltransferase | 0 | 2.45 | 2.26 |
| 2639 | Phosphoribosylaminoimidazole-succinocarboxamide synthase | 0 | 7.36 | 1.80 |
| 6314 | Phosphoadenylylsulfate reductase (thioredoxin) | 0 | 7.36 | 11.28 |
| 2240 | Phosphoribosyltransferase | 0 | 19.62 | 5.41 |
| 4194 | Phosphoribosylformylglycinamidine synthase II | 2.66 | 0 | 0 |
| 2553 | Adenylosuccinate synthetase | 2.66 | 0 | 2.26 |
| 3424 | Phosphomethylpyrimidine kinase | 2.66 | 0 | 0.90 |
| 6546 | Dihydroorotase | 2.66 | 2.45 | 0 |
| 2309 | Orotate phosphoribosyltransferase | 2.66 | 2.45 | 1.80 |
| 6261 | Bifunctional phosphoribosylaminoimidazolecarboxamide formyltransferase/IMP cyclohydrolase | 5.33 | 0 | 1.80 |
| 6333 | Adenylosuccinate lyase | 7.99 | 0 | 4.96 |
| Transcriptional regulation |  |  |  |  |
| RNA polymerase |  |  |  |  |
| 608 | RNA polymerase, sigma 70 subunit, RpoD subfamily | 0 | 0 | 0.45 |
| 1794 | DNA-directed RNA polymerase subunit omega | 0 | 0 | 3.61 |
| 2126 | RNA polymerase, sigma subunit, RpsC/SigC | 0 | 0 | 3.61 |
| 1436 | RNA polymerase sigma factor | 0 | 0 | 3.16 |
| 1465 | RpoD family RNA polymerase sigma factor | 0 | 0 | 0.45 |
| 1436 | RNA polymerase sigma factor | 0 | 0 | 3.16 |
| 1571 | DNA-directed RNA polymerase gamma chain | 5.33 | 0 | 4.96 |
| 2612 | DNA-directed RNA polymerase subunit alpha | 15.98 | 29.43 | 13.08 |
| 1572 | DNA-directed RNA polymerase beta' subunit | 37.30 | 2.45 | 3.16 |
| 1569 | DNA-directed RNA polymerase subunit beta | 39.96 | 2.45 | 3.61 |
| **Transcriptional regulator** |  |  |  |  |
| 3282 | SOS-response transcriptional repressor, LexA | 0 | 0 | 0.45 |
| 2666 | Transcriptional regulator, LysR family | 0 | 0 | 9.92 |
| 5981 | Transcriptional regulator, BolA protein family | 0 | 0 | 1.35 |
| 3539 | Signal recognition particle protein | 0 | 0 | 4.511 |
| 4517 | Two component transcriptional regulator | 0 | 2.45 | 0.90 |
| 1692 | Transcriptional regulator, GntR family | 0 | 4.90 | 0.45 |
| 3023 | Transcriptional regulator | 0 | 4.90 | 0.90 |
| 3487 | Transcriptional regulator, LysR family | 0 | 4.90 | 3.61 |
| 1692 | Transcriptional regulator, GntR family | 0 | 4.90 | 0.45 |
| 4132 | Anti-sigma-factor antagonist | 0 | 7.36 | 1.80 |
| 6455 | Response regulator receiver protein | 0 | 12.26 | 2.26 |
| 1420 | Two component LuxR family transcriptional regulator | 5.33 | 4.90 | 3.16 |
| 4375 | Putative transcriptional regulator | 7.99 | 0 | 0.45 |
| 1173 | Two component Transcriptional regulator, Winged helix family protein | 10.66 | 29.43 | 5.86 |
| 3590 | Two component LuxR family transcriptional regulator | 26.64 | 14.71 | 10.83 |
| 557 | AbrB family transcriptional regulator | 39.96 | 31.88 | 29.32 |
| **Protein synthesis** |  |  |  |  |
| **Amino acid synthesis** |  |  |  |  |
| 524 | Amidohydrolase 2 | 0 | 0 | 0.45 |
| 1031 | Amidohydrolase | 0 | 0 | 0.45 |
| 6700 | Aromatic amino acid beta-eliminating lyase/threonine aldolase | 0 | 4.9 | 0 |
| 985 | Diaminopimelate epimerase | 0 | 4.90 | 4.06 |
| 2185 | Acetylglutamate kinase | 0 | 9.81 | 0.90 |
| 2435 | Aspartate-semialdehyde dehydrogenase | 0 | 4.90 | 1.35 |
| 6314 | Phosphoadenylylsulfate reductase (thioredoxin) | 0 | 7.36 | 11.28 |
| 2446 | Saccharopine dehydrogenase | 7.99 | 0 | 0 |
| 3933 | Taurine catabolism dioxygenase TauD/TfdA | 23.98 | 0 | 0 |
| 3947 | Amidohydrolase 2 | 31.97 | 0 | 0 |
| **tRNA synthetase** |  |  |  |  |
| 260 | Aspartyl-glutamyl-tRNA amidotransferase subunit A | 0 | 0 | 4.06 |
| 5328 | Cysteinyl-tRNA synthetase | 0 | 0 | 0.45 |
| 39 | Glutamyl-tRNA reductase | 0 | 0 | 0.45 |
| 5530 | Isoleucyl-tRNA synthetase | 0 | 0 | 0.45 |
| 2962 | Leucyl-tRNA synthetase | 0 | 0 | 1.35 |
| 4316 | Methionyl-tRNA synthetase | 0 | 0 | 1.35 |
| 520 | Putative histidyl-tRNA synthetase 2 | 0 | 0 | 6.32 |
| 1748 | Seryl-tRNA synthetase | 0 | 2.45 | 2.71 |
| 1102 | Tryptophanyl-tRNA synthetase | 0 | 2.45 | 0 |
| 3220 | Tyrosyl-tRNA synthetase | 0 | 2.45 | 1.35 |
| 81 | Peptidyl-tRNA hydrolase | 0 | 4.90 | 4.06 |
| 3740 | Methionyl-tRNA formyltransferase | 0 | 7.36 | 4.06 |
| 4380 | Phenylalanyl-tRNA synthetase alpha subunit | 0 | 14.71 | 3.16 |
| 1850 | Glutamyl-tRNA(Gln) amidotransferase, B subunit | 2.66 | 0 | 5.86 |
| 2270 | Glycyl-tRNA synthetase alpha chain | 2.66 | 0 | 0 |
| 6531 | Histidyl-tRNA synthetase | 2.66 | 7.36 | 2.71 |
| 6635 | Arginyl-tRNA synthetase | 5.33 | 0 | 4.51 |
| 2649 | Glycyl and Arginyl tRNA synthetase | 7.99 | 0 | 10.83 |
| 4909 | Prolyl-tRNA synthetase | 7.99 | 0 | 4.96 |
| 3273 | Glutamyl-tRNA synthetase | 7.99 | 2.45 | 1.80 |
| 3210 | Lysyl-tRNA synthetase | 10.66 | 0 | 3.61 |
| 495 | Phenylalanyl-tRNA synthetase, beta subunit | 10.66 | 0 | 0 |
| 3786 | Valyl-tRNA synthetase | 10.66 | 0 | 2.26 |
| 817 | Threonyl-tRNA synthetase / Ser-tRNA(Thr) hydrolase | 13.32 | 0 | 0.45 |
| 4277 | Asparaginyl-tRNA synthetase | 15.98 | 0 | 0 |
| **Ribosomal protein** |  |  |  |  |
| 685 | 50S ribosomal protein L28 | 0 | 0 | 4.06 |
| 1006 | Ribosomal protein S21 | 0 | 0 | 8.57 |
| 1567 | Ribosomal protein S20 | 0 | 0 | 10.38 |
| 1713 | 30S ribosomal protein S18 | 0 | 0 | 4.06 |
| 1999 | Ribosomal protein L27 | 0 | 0 | 4.51 |
| 2591 | Ribosomal protein S19 | 0 | 0 | 13.98 |
| 2594 | LSU ribosomal protein L16P | 0 | 0 | 9.02 |
| 2595 | Ribosomal protein L29 | 0 | 0 | 6.77 |
| 2598 | Ribosomal protein L24 | 0 | 0 | 4.96 |
| 2603 | LSU ribosomal protein L18P | 0 | 0 | 3.61 |
| 2617 | 50S ribosomal protein L31 | 0 | 0 | 3.61 |
| 3791 | 30S ribosomal protein 3 | 0 | 0 | 2.71 |
| 4679 | 50S ribosomal protein L20 | 0 | 0 | 9.92 |
| 5266 | Ribosomal protein S15 | 0 | 0 | 3.16 |
| 2554 | 50S ribosomal protein L25 | 0 | 2.45 | 4.96 |
| 2592 | 50S ribosomal protein L22 | 0 | 2.45 | 7.22 |
| 2596 | 30S ribosomal protein S17 | 0 | 2.45 | 4.51 |
| 4086 | 30S ribosomal protein S6 | 0 | 2.45 | 4.06 |
| 2613 | 50S ribosomal protein L17 | 0 | 4.90 | 10.38 |
| 2973 | 50S ribosomal protein L9 | 0 | 4.90 | 11.28 |
| 6002 | Ribosomal protein L11 methyltransferase | 0 | 4.90 | 0 |
| 2588 | LSU ribosomal protein L4P | 0 | 12.26 | 10.83 |
| 6310 | Sigma 54 modulation protein/ribosomal protein S30EA | 0 | 19.62 | 18.50 |
| 1549 | 50S ribosomal protein L7/L12 | 2.66 | 0 | 22.56 |
| 2590 | 50S ribosomal protein L2 | 2.66 | 2.45 | 10.38 |
| 2609 | SSU ribosomal protein S13P | 2.66 | 2.45 | 13.08 |
| 5950 | 30S ribosomal protein S12 | 2.66 | 2.45 | 3.16 |
| 1543 | 50S ribosomal protein L19 | 2.66 | 4.90 | 3.61 |
| 2615 | LSU ribosomal protein L13P | 2.66 | 7.36 | 10.83 |
| 2589 | LSU ribosomal protein L23P | 2.66 | 9.81 | 7.67 |
| 2601 | 50S ribosomal protein L6 | 2.66 | 9.81 | 7.67 |
| 5954 | Ribosomal protein S10 | 2.66 | 12.26 | 11.73 |
| 1548 | 50S ribosomal protein L10 | 5.33 | 4.90 | 3.61 |
| 3538 | 30S ribosomal protein S16 | 5.33 | 4.90 | 12.18 |
| 5685 | 30S ribosomal protein S14 | 5.33 | 4.90 | 8.12 |
| 5951 | SSU ribosomal protein S7P | 5.33 | 4.90 | 20.75 |
| 2610 | SSU ribosomal protein S11P | 5.33 | 7.36 | 3.61 |
| 2616 | 30S ribosomal protein S9 | 5.33 | 7.36 | 8.12 |
| 2597 | LSU ribosomal protein L14P | 5.33 | 12.26 | 8.57 |
| 2586 | LSU ribosomal protein L3P | 5.33 | 19.62 | 7.67 |
| 4920 | SSU ribosomal protein S4P | 7.99 | 2.45 | 4.96 |
| 2605 | 50S ribosomal protein L15 | 10.66 | 14.71 | 17.14 |
| 1547 | 50S ribosomal protein L1 | 10.66 | 26.97 | 21.65 |
| 2600 | 30S ribosomal protein S8 | 13.32 | 9.81 | 8.12 |
| 2604 | 30S ribosomal protein S5 | 13.32 | 17.17 | 24.36 |
| 2599 | 50S ribosomal protein L5 | 13.32 | 22.07 | 21.65 |
| 1546 | 50S ribosomal protein L11 | 15.98 | 22.07 | 16.24 |
| 2593 | SSU ribosomal protein S3P | 18.65 | 14.71 | 9.47 |
| 4068 | SSU ribosomal protein S2P | 21.31 | 12.26 | 10.83 |
| 5643 | 30S ribosomal protein S1 | 26.64 | 26.97 | 4.96 |
| Translation initiation factor |  |  |  |  |
| 2608 | Bacterial translation initiation factor 1 (bIF-1) | 0 | 0 | 5.86 |
| 5811 | Translation initiation factor IF-3 | 0 | 0 | 5.41 |
| 3766 | Translation initiation factor IF-2 | 13.32 | 0 | 2.71 |
| 5952 | Translation elongation factor 2 (EF-2/EF-G) | 50.62 | 7.36 | 12.63 |
| 5953 | Translation elongation factor 1A (EF-1A/EF-Tu) | 175.82 | 193.72 | 72.63 |

**ORF#**: open reading frame # were given by
http://scorpius.ucdavis.edu/gmod/cgibin/site/anabaena02?page=gblast

DNA-directed RNA polymerase subunits alpha-, beta, beta’ and gamma were abundant in akinetes. However some proteins required for transcription (like RNA polymerase sigma factor) and translation (like signal recognition particle protein, some tRNA and ribosomal proteins) were only found in heterocysts and vegetative cells (Table 2), suggesting a very active transcription and translation occurred in heterocysts and vegetative cells. Thirteen out of 25 tRNA synthetases and 23 out of 50 ribosomal proteins were absent in akinetes. Most transcriptional regulators were also undetectable in akinetes. These data imply that akinetes retain a less active transcription machinery and a very weak translational capability.

The proteomics data showed that amidohydrolase 2 (ORF: 3947) and taurine catabolism dioxygenase TauD/TfdA (ORF: 3933) were the third and fifth most abundant akinete-specific proteins, while the other two amidohydrolases were vegetative cell-specific (Table 2). TauD/TfdA, which can degrade taurine and be a source of sulfur (Shen et al., 2007), was found in akinetes as well. However, phosphoadenylylsulfate reductase, involved in sulfur (Wang et al., 2004) and pyrimidine metabolism (http://www.genome.jp/kegg-bin/show_pathway?ec00240 +1.8.1.9), was absent in akinetes. Certain enzymes involved in amino acid metabolism were only found in heterocysts and/or vegetative cells, such as arginine (Arg) biosynthetic enzyme, acetylglutamate kinase (Ramon-Maiques et al., 2002), and lysine biosynthetic enzyme diaminopimelate epimerase (Hor et al., 2013). Interestingly, saccharopine dehydrogenase required for lysine degradation (Serrano et al., 2012) was only present in akinetes.

### Cell division

Heterocysts, terminally differentiated N_2_-fixing cells, do not divide and need not pass DNA information to the next generation, which is consistent with the absence of key DNA replication enzymes (DNA polymerase) in heterocysts. Similarly, no DNA polymerases were detected in akinetes, suggesting that akinetes are not dividing either. Akinetes of *A. cylindrica* store twice as much DNA and 10-fold more protein than vegetative cells (Simon, 1977), preparing them for germination when environmental conditions become favorable. Surprisingly, septum formation protein Maf (Briley et al., 2011), participating in cell division in *Bacillus subtilis*, was found to be heterocyst-specific. Cell division protein FtsZ (Bi and Lutkenhaus, 1990) and septum site-determining protein MinD (Maurya et al., 2016) were more abundant in heterocysts and akinetes than in vegetative cells (Table 3). We speculated that, instead of involvement in cell division, these proteins might be critical in maintaining septum homeostasis among heterocysts, akinetes, and vegetative cell.

**TABLE 3.**
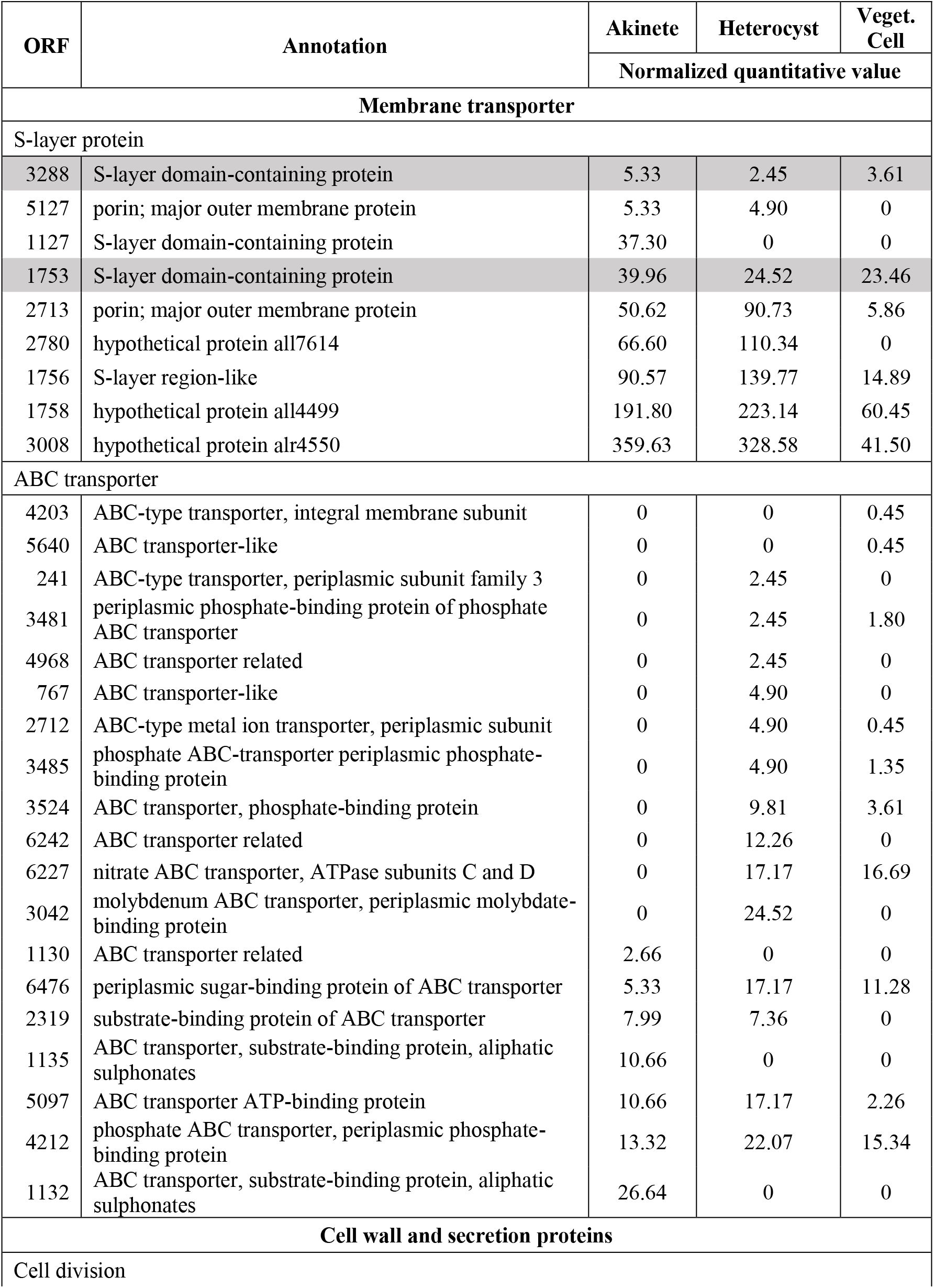
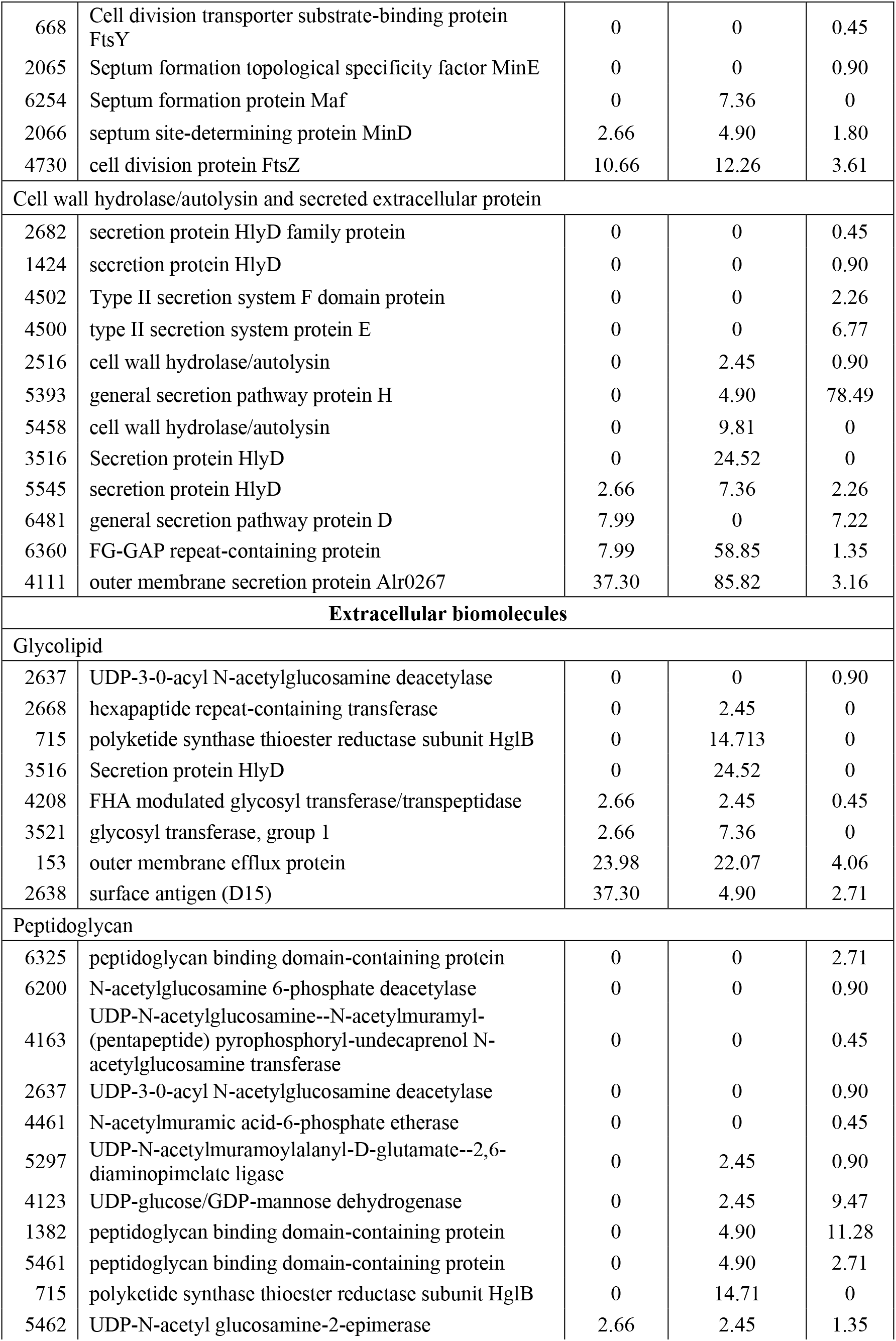
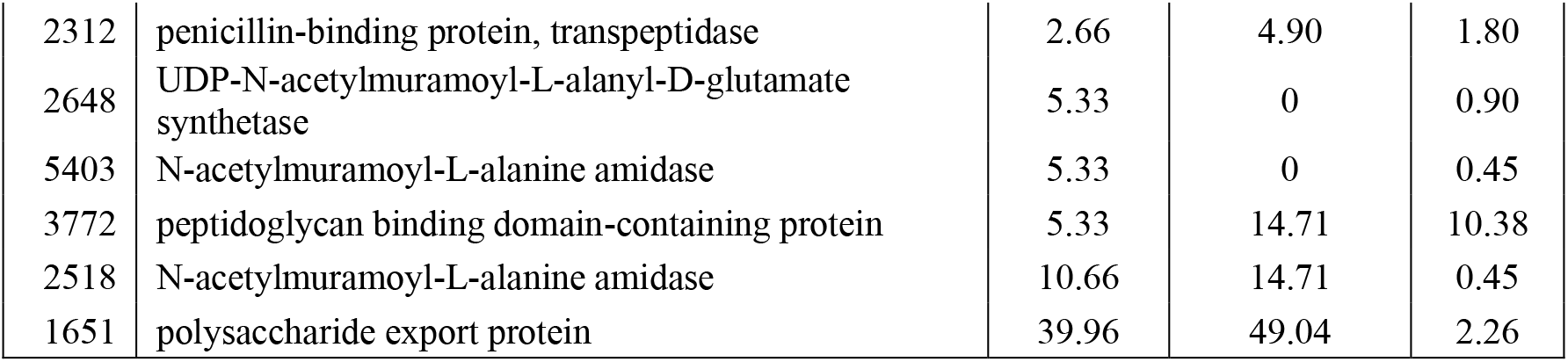
Proteins involved in synthesizing and transporting polysaccharides and peptidoglycan in akinetes, heterocysts, and vegetative cells (low similarity of the S-layer domain containing proteins are highlighted in grey)

| ORF | Annotation | Akinete | Heterocyst | Veget. Cell |
| --- | --- | --- | --- | --- |
|  |  | Normalized quantitative value |  |  |
| Membrane transporter |  |  |  |  |
| S-layer protein |  |  |  |  |
| 3288 | S-layer domain-containing protein | 5.33 | 2.45 | 3.61 |
| 5127 | porin; major outer membrane protein | 5.33 | 4.90 | 0 |
| 1127 | S-layer domain-containing protein | 37.30 | 0 | 0 |
| 1753 | S-layer domain-containing protein | 39.96 | 24.52 | 23.46 |
| 2713 | porin; major outer membrane protein | 50.62 | 90.73 | 5.86 |
| 2780 | hypothetical protein all7614 | 66.60 | 110.34 | 0 |
| 1756 | S-layer region-like | 90.57 | 139.77 | 14.89 |
| 1758 | hypothetical protein all4499 | 191.80 | 223.14 | 60.45 |
| 3008 | hypothetical protein alr4550 | 359.63 | 328.58 | 41.50 |
| ABC transporter |  |  |  |  |
| 4203 | ABC-type transporter, integral membrane subunit | 0 | 0 | 0.45 |
| 5640 | ABC transporter-like | 0 | 0 | 0.45 |
| 241 | ABC-type transporter, periplasmic subunit family 3 | 0 | 2.45 | 0 |
| 3481 | periplasmic phosphate-binding protein of phosphate ABC transporter | 0 | 2.45 | 1.80 |
| 4968 | ABC transporter related | 0 | 2.45 | 0 |
| 767 | ABC transporter-like | 0 | 4.90 | 0 |
| 2712 | ABC-type metal ion transporter, periplasmic subunit | 0 | 4.90 | 0.45 |
| 3485 | phosphate ABC-transporter periplasmic phosphate-binding protein | 0 | 4.90 | 1.35 |
| 3524 | ABC transporter, phosphate-binding protein | 0 | 9.81 | 3.61 |
| 6242 | ABC transporter related | 0 | 12.26 | 0 |
| 6227 | nitrate ABC transporter, ATPase subunits C and D | 0 | 17.17 | 16.69 |
| 3042 | molybdenum ABC transporter, periplasmic molybdate-binding protein | 0 | 24.52 | 0 |
| 1130 | ABC transporter related | 2.66 | 0 | 0 |
| 6476 | periplasmic sugar-binding protein of ABC transporter | 5.33 | 17.17 | 11.28 |
| 2319 | substrate-binding protein of ABC transporter | 7.99 | 7.36 | 0 |
| 1135 | ABC transporter, substrate-binding protein, aliphatic sulphonates | 10.66 | 0 | 0 |
| 5097 | ABC transporter ATP-binding protein | 10.66 | 17.17 | 2.26 |
| 4212 | phosphate ABC transporter, periplasmic phosphate-binding protein | 13.32 | 22.07 | 15.34 |
| 1132 | ABC transporter, substrate-binding protein, aliphatic sulphonates | 26.64 | 0 | 0 |
| Cell wall and secretion proteins |  |  |  |  |
| Cell division |  |  |  |  |
| 668 | Cell division transporter substrate-binding protein FtsY | 0 | 0 | 0.45 |
| 2065 | Septum formation topological specificity factor MinE | 0 | 0 | 0.90 |
| 6254 | Septum formation protein Maf | 0 | 7.36 | 0 |
| 2066 | septum site-determining protein MinD | 2.66 | 4.90 | 1.80 |
| 4730 | cell division protein FtsZ | 10.66 | 12.26 | 3.61 |
| Cell wall hydrolase/autolysin and secreted extracellular protein |  |  |  |  |
| 2682 | secretion protein HlyD family protein | 0 | 0 | 0.45 |
| 1424 | secretion protein HlyD | 0 | 0 | 0.90 |
| 4502 | Type II secretion system F domain protein | 0 | 0 | 2.26 |
| 4500 | type II secretion system protein E | 0 | 0 | 6.77 |
| 2516 | cell wall hydrolase/autolysin | 0 | 2.45 | 0.90 |
| 5393 | general secretion pathway protein H | 0 | 4.90 | 78.49 |
| 5458 | cell wall hydrolase/autolysin | 0 | 9.81 | 0 |
| 3516 | Secretion protein HlyD | 0 | 24.52 | 0 |
| 5545 | secretion protein HlyD | 2.66 | 7.36 | 2.26 |
| 6481 | general secretion pathway protein D | 7.99 | 0 | 7.22 |
| 6360 | FG-GAP repeat-containing protein | 7.99 | 58.85 | 1.35 |
| 4111 | outer membrane secretion protein Alr0267 | 37.30 | 85.82 | 3.16 |
| Extracellular biomolecules |  |  |  |  |
| Glycolipid |  |  |  |  |
| 2637 | UDP-3-0-acyl N-acetylglucosamine deacetylase | 0 | 0 | 0.90 |
| 2668 | hexapaptide repeat-containing transferase | 0 | 2.45 | 0 |
| 715 | polyketide synthase thioester reductase subunit HglB | 0 | 14.713 | 0 |
| 3516 | Secretion protein HlyD | 0 | 24.52 | 0 |
| 4208 | FHA modulated glycosyl transferase/transpeptidase | 2.66 | 2.45 | 0.45 |
| 3521 | glycosyl transferase, group 1 | 2.66 | 7.36 | 0 |
| 153 | outer membrane efflux protein | 23.98 | 22.07 | 4.06 |
| 2638 | surface antigen (D15) | 37.30 | 4.90 | 2.71 |
| Peptidoglycan |  |  |  |  |
| 6325 | peptidoglycan binding domain-containing protein | 0 | 0 | 2.71 |
| 6200 | N-acetylglucosamine 6-phosphate deacetylase | 0 | 0 | 0.90 |
| 4163 | UDP-N-acetylglucosamine--N-acetylmuramyl-(pentapeptide) pyrophosphoryl-undecaprenol N-acetylglucosamine transferase | 0 | 0 | 0.45 |
| 2637 | UDP-3-0-acyl N-acetylglucosamine deacetylase | 0 | 0 | 0.90 |
| 4461 | N-acetylmuramic acid-6-phosphate etherase | 0 | 0 | 0.45 |
| 5297 | UDP-N-acetylmuramoylalanyl-D-glutamate--2,6-diaminopimelate ligase | 0 | 2.45 | 0.90 |
| 4123 | UDP-glucose/GDP-mannose dehydrogenase | 0 | 2.45 | 9.47 |
| 1382 | peptidoglycan binding domain-containing protein | 0 | 4.90 | 11.28 |
| 5461 | peptidoglycan binding domain-containing protein | 0 | 4.90 | 2.71 |
| 715 | polyketide synthase thioester reductase subunit HglB | 0 | 14.71 | 0 |
| 5462 | UDP-N-acetyl glucosamine-2-epimerase | 2.66 | 2.45 | 1.35 |
| *Proteomics of three cell types* |  |  |  |  |
| 2312 | penicillin-binding protein, transpeptidase | 2.66 | 4.90 | 1.80 |
| 2648 | UDP-N-acetylmuramoyl-L-alanyl-D-glutamate synthetase | 5.33 | 0 | 0.90 |
| 5403 | N-acetylmuramoyl-L-alanine amidase | 5.33 | 0 | 0.45 |
| 3772 | peptidoglycan binding domain-containing protein | 5.33 | 14.71 | 10.38 |
| 2518 | N-acetylmuramoyl-L-alanine amidase | 10.66 | 14.71 | 0.45 |
| 1651 | polysaccharide export protein | 39.96 | 49.04 | 2.26 |

### Heterocyst-specific envelope glycolipid and lipopolysaccharide lipid A

Cyanobacterial heterocysts provide a micro-oxic environment to support the oxygen-labile nitrogenase fixing N_2_ in an oxic milieu. The heterocyst glycolipid (HGL) layer is an important part of the system for maintaining a micro-oxic environment in heterocysts (Murry and Wolk, 1989). Our proteomics study identified multiple heterocyst-specific proteins required for synthesis, export, and deposition of envelope polysaccharides and glycolipids (Table 3). For instance, we found polyketide synthase thioester reductase subunit HglB (Fan et al., 2005), an enzyme for synthesizing glycolipid aglycones. DevA and DevB are two components of DevBCA exporter (Fiedler et al., 1998) necessary for the formation of the laminated layer of heterocysts (Zhou and Wolk, 2003). Hexapeptide repeat-containing transferase is a sugar transferase which might play a critical role in synthesizing different sugars from the fixed carbon source provided by adjacent vegetative cells (Vaara, 1992). Furthermore, glycosyl transferase (HglT, ORF: 3521) required to glycosylate the glycolipid aglycone (Awai and Wolk, 2007) was present only in heterocysts and akinetes (Table 3). ORF: 2637 and ORF: 2638, orthologs of LpcC and Omp85 involved in lipopolysaccharide lipid A biosynthesis to form a permeability barrier at the outer membrane (Nicolaisen et al., 2009), had different distribution, with high abundance of Omp85 in akinetes, supporting the hypothesis that the lipopolysaccharide layer plays an important role in increasing stress tolerance of akinetes in *A. cylindrica*.

### S-layer proteins and ATP-binding cassette (ABC) transporter

The *A. cylindrica* genome encodes seven S-layer domain-containing proteins (Supplementary Figure S2) and two S-layer like proteins. S-layer proteins can be self-assembled to form an array on the surface of the cell (Smarda et al., 2002). They have multiple functions, including the maintenance of cell integrity, a permeability barrier, pathogenesis, and immune response (Gerbino et al., 2015). Our LC-MS/MS identified all nine S-layer proteins (Table 3). Most S-layer proteins, e.g., all4499 and alr4550 (Oliveira et al., 2015), along with other extracellular proteins, such as FG-GAP repeat-containing protein HesF (Oliveira et al., 2015) (Table 3), have been identified as exoproteins. The abundance of these nine S-layer proteins was different, with one S-layer protein (ORF: 1127) unique to akinetes, and two other S-layer proteins (ORF: 5127 and ORF: 2780) absent in vegetative cells (Table 3), implying unique functionality associated to different cell types.

### Polysaccharide and Peptidoglycan in cyanobacterial cell wall

We identified a total of 17 proteins involved in peptidoglycan and lipopolysaccharides (LPS) formation, among them, 7, 10, and 16 proteins related to peptidoglycan and lipopolysaccharides were found in akinetes, heterocysts and vegetative cells, respectively (Table 3). Most cyanobacteria have an additional polysaccharide layer in the cell envelope (Cardemil and Wolk, 1976; 1979). S-layer proteins are anchored to the cell surface through non-covalent interactions with cell surface structures, usually containing LPS (Gandham et al., 2012). UDP-glucose/GDP-mannose dehydrogenase, which takes part in the synthesis of LPSs (Muszynski et al., 2011) was found in heterocysts and vegetative cells, but absent in akinetes. Notably, orthologs of Alr2887 (ORF: 153) involved in heterocyst-specific glycolipid export and All4388 (ORF: 1651) involved in heterocyst envelope polysaccharide deposition (Maldener et al., 2003) were shown more abundant in akinetes and heterocysts (Table 3), suggesting a role in envelope formation of heterocysts and akinetes.

### Glycogen serves as a form of energy storage

Glycogen is a multibranched polymer of glucose serving as the major carbon storage in cyanobacteria (Diaz-Troya et al., 2014). Glycogen biosynthesis is coupled to photosynthesis, and its conversion into glucose in the dark is necessary to maintain cell metabolism. ADP-glucose pyrophosphorylase (AGP) and glycogen synthase are required for synthesis of glycogen. Interestingly, glycogen synthase was highly abundant in akinetes and rare in vegetative cells. Neither akinetes nor heterocysts contained the five proteins involved in glycogen degradation (Table 4).

**TABLE 4.** Proteins involved in metabolism of cyanophycin, glycogen, and sucrose in akinetes, heterocysts, and vegetative cells (enzymes responsible for anabolism are highlighted in grey)

| ORF | Annotation | Akinete | Heterocyst | Veget. Cell |
| --- | --- | --- | --- | --- |
|  |  | Normalized quantitative value |  |  |
| Cyanophycin/arginine |  |  |  |  |
| 1510 | cyanophycin synthetase | 2.66 | 0 | 0 |
| 1272 | Cyanophycinase, Serine peptidase, MEROPS family S51 | 7.99 | 4.90 | 0.45 |
| 3212 | putative cyanophycinase | 0 | 12.26 | 0 |
| 1511 | cyanophycinase | 0 | 9.81 | 2.26 |
| 4256 | isoaspartyl dipeptidase, peptidase T2, asparaginase 2 | 0 | 0 | 4.96 |
| 2185 | acetylglutamate kinase | 0 | 9.81 | 0.90 |
| 1480 | N-acetyl-gamma-glutamyl-phosphate reductase | 10.66 | 19.62 | 5.86 |
| 3400 | Nitrogen regulatory protein P-II (GlnB, GlnK) | 5.33 | 29.43 | 5.86 |
| Sucrose/trehalose |  |  |  |  |
| 4634 | sucrose synthase SuS-B | 21.31 | 0 | 0 |
| 3602 | sucrose synthase SuS-A | 15.98 | 0 | 0 |
| 4573 | neutral invertase InvB | 0 | 0 | 0.45 |
| 3842 | sucrose phosphatase SPP | 0 | 7.36 | 1.35 |
| 238 | malto-oligosyltrehalose synthase | 5.33 | 0 | 0 |
| Glycogen |  |  |  |  |
| 1048 | glycogen synthase | 21.31 | 12.26 | 1.35 |
| 2144 | phosphoglucomutase/phosphomannomutase | 13.32 | 12.26 | 20.30 |
| 3396 | phosphoglucomutase/phosphomannomutase alpha/beta/subunit | 5.33 | 2.45 | 1.80 |
| 6292 | 1,4-alpha-glucan-branching enzyme | 2.66 | 0 | 0.45 |
| 5134 | glycogen debranching enzyme GlgX | 0 | 0 | 0.45 |
| 341 | Phosphoglycerate/bisphosphoglycerate mutase | 0 | 0 | 0.45 |
| 5891 | Phosphoglycerate mutase | 0 | 0 | 0.45 |
| 6104 | phosphoglycerate mutase, 2,3-bisphosphoglycerate-independent | 0 | 0 | 3.61 |
| 5419 | phosphoglycerate mutase | 0 | 0 | 0.90 |

### Cyanophycin and β-aspartyl-arginine

Cyanophycin (CpG), or multi-L-arginyl-poly-L-aspartic acid granule polypeptide, is a non-ribosomally produced amino acid polymer composed of an aspartic acid (Asp) backbone and Arg side groups. In heterocysts, nitrogenase converts N_2_ to ammonia and then forms glutamine (Gln). Gln can serve as ammonium donor for synthesis of Asp by aspartate aminotransferase, also known as glutamic oxaloacetic transaminase (Xu et al., 2015). Gln is also the precursor for biosynthesis of Arg and proteins involved in Arg biosynthesis, and it was found in high amount in heterocysts, such as acetylglutamate kinase (Huang et al., 2015;Minaeva et al., 2015), ArgL (ORF: 1480) (Leganes et al., 1998), and nitrogen regulatory protein P-II GlnB (ORF: 3400) (Llacer et al., 2007; Paz-Yepes et al., 2009) (Table 4). Asp and Arg are further condensed by cyanophycin synthetase into CpG (Ziegler et al., 1998). This nitrogen storage molecule can be degraded by cyanophycinase (Picossi et al., 2004) to produce β-aspartyl-arginine. Cyanophycin synthetase was below the limit of detection in heterocysts, but all three putative cyanophycinases in the *A. cylindrica* genome were present in high amounts (Table 4), supporting a previous finding that cyanophycinase activity is high in heterocysts (Gupta and Carr, 1981). Asp and Arg can also be transported into akinetes for further condensation into CpG by cyanophycin synthetase (ORF: 1510) and stored. CpG can be degraded by cyanophycinase (ORF: 1272) (Table 4) to support growth of other cells in the filament and/or germination in a favorable environment.

### Sucrose as a reducing power for N_2_ fixation and compatible solute

Sucrose, a universal vehicle of reduced carbon in plants, appears to have a similar role within the diazotrophic cyanobacterial filament (Kolman et al., 2015). Sucrose synthesized by sucrose phosphate synthase and sucrose phosphate phosphatase (SPP) is believed to occur in *Anabaena* strains (Cumino et al., 2002). Sucrose is then transported into heterocysts (Juttner, 1983) and further hydrolyzed by a specific invertase (InvB) (Lopez-Igual et al., 2010;Vargas et al., 2011). The bidirectional enzyme sucrose synthase SuS-A, on the other hand, exhibited optimal activity at pH 7.5-8.2 in the sucrose-synthesis direction and at pH 5.9-6.5 in the reverse direction (Porchia et al., 1999). Our proteomic data identified SPP (ORF: 3842) (Cumino et al., 2001) present in heterocysts and vegetative cells, but not in akinetes. More strikingly, we detected both SuS-A (ORF: 3602) and SuS-B (ORF: 4634) in high amount, but no invertase in akinetes. Invertase in heterocysts was below the limit of detection in heterocysts. We speculated that the high amount of sucrose synthase present in akinetes might be involved in breaking down sucrose transported from vegetative cells for synthesizing reserve glycogen (Perez et al., 2016), polysaccharides to build akinete envelope, and/or for synthesizing trehalose as an osmoprotectant (Sakamoto et al., 2009) by akinete-specific malto-oligosyltrehalose synthase (ORF: 238) orthologous to All0167 (Higo et al., 2006). Trehalose may play a role in long-term survival of akinetes under dry conditions.

## Discussion

Some filamentous cyanobacteria can differentiate nitrogen-fixing cells called heterocysts. Normally 2 ~ 10% of vegetative cells develop into heterocysts. In *A. cylindrica*, vegetative cells adjoining heterocysts develop into akinetes (Figure 1A). The vegetative cells capture sunlight energy to fix CO_2_ and heterocysts carry out solar-powered N_2_-fixation. Although akinetes are known as spore-like structures for survival under unfavorable condition, our proteomic data indicate that akinetes may also play an active role during filamentous growth. Based on the distribution of cyanophycin, glycogen and sucrose biosynthesis-related proteins, a putative network for fixed nitrogen and carbohydrate among Heterocysts, Akinetes and Vegetative cells, or designated HAVe model, is proposed for *A. cylindrica* (Figure 3). This is the first comprehensive comparison of proteins of akinetes, heterocysts and vegetative cells of *A. cylindrica*. These findings support new insight into the metabolic differences and increase our understanding of the roles played by these three very different but adjacent cells. The distinct distribution of FOX proteins, PS I & II proteins, and AcaK43 in heterocysts, vegetative cells, and akinetes, respectively, is consistent with previous findings, supporting the reliability of our proteomic data. Only the RuBisCO results (Table S2) are inconsistent with the previous observations. We observed high abundance of RuBisCO large and small subunits, and some carboxysomal microcompartment proteins (CcmN, CcmM, ORFs: 2671-2672) in all three cell types (Table S2, (Cameron et al., 2013)). However, Cossar et al. reported that RuBisCO protein was undetectable in mature heterocysts of *A. cylindrica* (Cossar et al., 1985). Several lines of evidence from *Anabaena* strain PCC 7120 have shown that promoter activity of RuBisCO was barely detected in heterocysts using P*rbcLS*-luxAB as a reporter (Elhai and Wolk, 1990), and RuBisCO large and small subunit transcripts were not detected in heterocysts by *in situ* hybridization (Madan and Nierzwicki-Bauer, 1993). Whether RuBisCO plays a role in both heterocysts and akinetes of *A. cylindrica* remains to be further investigated.

**FIGURE 3.**
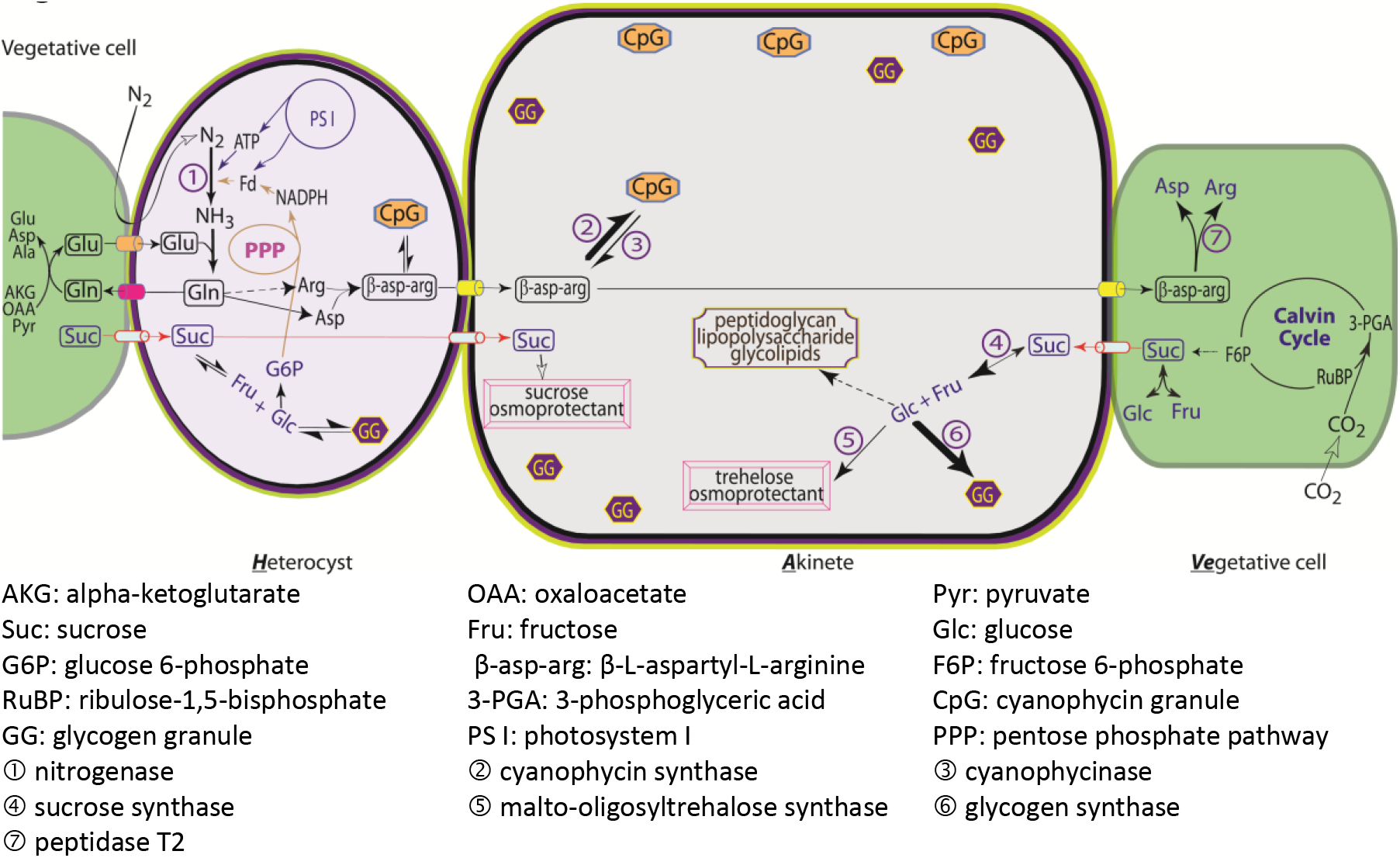
The HAVe (Heterocysts, Akinetes and Vegetative cells) model suggesting metabolic networks of cyanophycin and carbohydrates among heterocysts, akinetes, and vegetative cells.

The proteome of vegetative cells confirmed much of what is known about these workhorses. The large complement of PS I and PS II proteins supported active photosynthesis while RuBisCO, carboxysomal proteins, and other enzymes of the Calvin cycle supported carbon fixation. The glucose and fructose produced is likely synthesized into sucrose in vegetative cells and then supplied to the adjoining akinetes and heterocysts as primary energy and carbon source (Figure 3). The HAVe model was supported by the findings of carbohydrate related proteins in this proteomics study. In vegetative cells, fructose 6-phosphate is generated via the Calvin cycle during photosynthesis, which is then converted to sucrose by sucrose-phosphate synthase (SPS) and sucrose-phosphate phosphatase (SPP) (Cumino et al., 2002). Sucrose can be broken down by invertase in vegetative cells, or transported to akinetes where sucrose is cleaved into glucose and fructose, serving as building blocks for other carbohydrate biosynthesis, e.g., peptidoglycan, lipopolysaccharide, and glycolipid as envelope materials; glycogen storage molecules; and/or trehalose osmoprotectant. The paucity of FOX proteins along with key components of nitrogenase such as NifD, NifN and NifU not detected supported absence of nitrogen fixation. The FOX protein, Histone-like DNA binding protein HanA was most abundant in vegetative cells, consistent with observations that a strong HanA-GFP fluorescent signal co-localized with DNA in vegetative cells (Lu et al., 2014). A HanA mutant exhibited slow growth, altered pigmentation, and inability to differentiate heterocysts (Khudyakov and Wolk, 1996). FOX proteins unique to vegetative cells included trace amount of HepN (Lechno-Yossef et al., 2006), InvA, FraG, PrpI, NifU-like, DevR (Campbell et al., 1996), and H6L region containing protein (ORF: 2881). Vegetative cells obtain fixed nitrogen from either heterocysts or adjoining akinetes in the form of β-aspartyl-arginine. β-aspartyl-arginine is further degraded into Asp and Arg by isoaspartyl dipeptidase (ORF: 4256) in vegetative cells (Table 4, (Burnat et al., 2014)). Asp and Arg in turn serve as precursors for the biosynthesis of other amino acids and nucleotides, the building blocks for DNA, RNA, and protein biosynthesis. Vegetative cells contained abundant enzymes for nucleotide and amino acid biosynthesis. Forty-nine out of 50 ribosomal proteins and translation factors were found in vegetative cells as well. These data suggest that DNA, RNA, and protein biosynthesis occurs actively in vegetative cells to maintain their cellular function and cell division.

The heterocyst proteome supported what is known about these specialized cells, but also indicated some novel functions. Heterocysts contained all the proteins required for nitrogen fixation, including several proteins absent in akinetes and vegetative cells (Table 1, S1). These included nitrogenase molybdenum-iron protein NifN (Hu et al., 2010), the Fe-S cluster scaffold protein NifU that facilitates functional expression of nitrogenase in heterocysts (Nomata et al., 2015), and DevA required for heterocyst maturation (Maldener et al., 1994). Nitrogenase iron protein NifH (Mevarech et al., 1980) had high abundance in heterocysts, but was barely detected in vegetative cells and undetectable in akinetes (Table 1). Thus, the distribution of both Nif and Fox proteins indicated that N_2_-fixation only occurred in heterocysts. Ammonia produced by nitrogenase is incorporated into glutamine (Gln), serving as ammonia donor to Asp and Arg. Asp and Arg are condensed by cyanophycin synthetase into cyanophycin in heterocysts (Burnat et al., 2014). The high levels of three cyanophycinases in heterocysts (Table 4) indicate that the bulk of fixed nitrogen is then available as β-aspartyl-arginine, a nitrogen vehicle to be transferred intercellularly to be either hydrolyzed into Asp and Arg in the vegetative cells, or condensed into storage cyanophycin granule by cyanophycin synthetase in adjoining akinetes. All but three photosystem proteins occurred in heterocysts. The abundance of several PS I and PS II proteins implied at least partial functioning of PS I and PS II (Table 1). Generation of oxygen (O_2_) through PS II runs counter to the reductive process of nitrogenase. Nitrogenase is very sensitive to oxygen (O_2_), so the heterocysts must create a micro-oxic environment. Cytochrome C oxidase subunit II is the last enzyme in the respiratory electron transport chain. Valladares et al. showed that Cox2 and Cox3 transcription was up-regulated in heterocysts after nitrogen step-down in an NtcA- and HetR-dependent manner, and inactivation of both coxB2 and coxA3 results in the inability of *Anabaena* sp. PCC 7120 to grow diazotrophically under aerobic conditions (Valladares et al., 2003). Consistent with their observation, CoxB3 was found in akinetes and heterocysts, and CoxB2 was only found in heterocysts (Table 1). Taken together, as cytochrome C oxidase has high affinity to oxygen, it may play a role of consuming residual oxygen in heterocysts, and keeping nitrogenase in its active state. The high abundance of RuBisCO in heterocysts may play a role in removing residual oxygen by oxidizing Ribulose 1,5-bisphosphate into 3-PGA and 2-phosphoglycolic acid (Eisenhut et al., 2008). Flavodiiron protein Flv3B (ORF: 1739, homolog of all0178) was identified to be abundant in heterocysts (Table S2), which may also be responsible for light-induced O_2_ uptake in heterocysts to protect nitrogenase activity (Ermakova et al., 2014).

The key DNA replication proteins (DNA polymerases, SSBP) were not detected in heterocysts, consistent with terminal nature of the cells (Table 2). However, the presence of gyrase and helicase might play an important role in DNA rearrangement observed in heterocysts (Golden et al., 1985). Like vegetative cells, heterocysts contained a broad spectrum of proteins involved with transcription and translation. Heterocysts are encased in a thick envelope to supply a micro-oxic environment for protection of nitrogenase. Proteomic data indicated a number of heterocyst-specific proteins for synthesis, export, and external assembly of envelope polysaccharides and glycolipids (Table 3). Heterocysts were also decorated with six of the seven S-layer secretion proteins in Gram-negative species. Secretion relies on specific ATP-biding cassette (ABC) transporters and an outer membrane pores (Awram and Smit, 1998; Kawai et al., 1998). Heterocysts had several more ABC transporters (14) than did akinetes (7) or vegetative cells (10) (Table 3). The differential compositions of S-layer proteins and ABC transporters in the three cell types may contribute to the differences in cell envelope structure, including the greater resistance to cell disruption of heterocysts and also akinetes.

Akinetes appear to play a role as nitrogen and carbon storage cum transfer unit in filaments of *A. cylindrica* (Figure 3). By this model fixed carbon enters into akinetes from vegetative cells and is converted to glycogen by glycogen synthase, or into trehalose for osmoprotection during the survival stage. Heterocysts flanked by akinetes on both sides would then obtain carbon for energy via akinetes. Similarly akinetes receive β-aspartyl-arginine from heterocysts. This dipeptide is then either converted to cyanophycin for temporary or long-term storage, or transferred to the adjoining vegetative cells to support the growing chain. Our proteomic data also indicated that akinetes have less active transcriptional and translational machinery. Importantly, proteomic data indicated a cell envelope that was different to those of vegetative cells or heterocysts. Akinetes were decorated with all seven S-layer proteins detected. The suite of peptidoglycan synthesizing machinery and cell wall hydrolases differed (Table 2), as did the complement of membrane transporters and enzymes involved in polysaccharide structures. It is worthy to note that ORF2780 protein, homologous to carbohydrate-selective porins (OprB), functions as a sugar porin responsible for the optimal uptake of both fructose and glucose in Nostoc punctiforme ATCC 29133 (Ekman et al., 2013). The distinct distribution of OprB in akinetes and heterocysts at high abundance suggests a role in sugar uptake in these differentiated cells, consistent with the previous observation of carbon movement from vegetative cells to heterocysts of *A. cylindrica* (Wolk, 1968), which might also be true of carbon movement from vegetative cells to akinetes.

Akinetes have been viewed as spore like cells with the role of species survival under drought conditions. Their location between nitrogen-fixing heterocysts and carbon-fixing vegetative cells, combined with high levels of cyanophycin synthetase, cyanophycinase, sucrose synthetase, and glycogen synthetase suggests a critical role for akinetes during growth of *A. cylindrica* as demonstrated by the HAVe model. The role of the various genes and their regulation, as well as metabolite exchange among akinetes and their adjoining heterocysts and vegetative cells will need to be investigated in future work.

## Author Contributions

YQ, LG, and RZ designed the work. YQ, LG, MH, DW and RZ performed the experiments. YQ, LG, VB, DW, and RZ analyzed the proteomic data and drafted the manuscript. YQ, LG, VB, DW, MH and RZ revised the manuscript and responsible for final approval of the version to be published. All authors agree to be accountable for the content of the work.

## ACKNOWLEDGEMENTS

YQ was supported by the South Dakota Agricultural Experiment Station. The authors would like to acknowledge use of the South Dakota State University Functional Genomics Core Facility supported in part by NSF/EPSCoR Grant No. 0091948 and by the State of South Dakota

## SUPPLEMENTARY MATERIAL

The Supplementary Material for this article includes Figure S1 & S2, Table S1 & S2, and Supplemental information for LC-MS/MS data analyses.

